# Evolution-inspired dissection of caspase activities enables the redesign of caspase-4 into an LPS sensing interleukin-1 converting enzyme

**DOI:** 10.1101/2020.12.07.413732

**Authors:** Pascal Devant, Anh Cao, Jonathan C. Kagan

**Affiliations:** Division of Gastroenterology, Boston Children’s Hospital and Harvard Medical School, 300 Longwood Avenue, Boston, MA 02115, USA

## Abstract

Innate immune signaling pathways comprise multiple proteins that promote inflammation. This multistep means of information transfer suggests that complexity is a prerequisite for pathway design. Herein, we examined this possibility by studying caspases that regulate inflammasome-dependent inflammation. Several caspases differ in their ability to recognize bacterial LPS and cleave interleukin-1β (IL-1β). No caspase is known to contain both activities, yet distinct caspases with complementary activities bookend an LPS-induced pathway to IL-1β cleavage. Using unique caspases present in carnivorans as a guide, we identified molecular determinants of IL-1β cleavage specificity within caspase-1. This knowledge enabled the redesign of human caspase-4 to operate as a one-protein signaling pathway, which intrinsically links LPS detection to IL-1β cleavage and release, independent of inflammasomes. Strikingly, cat caspase-4 displays the activities of redesigned human caspase-4. These findings illustrate natural signaling pathway diversity and highlight how multistep innate immune pathways can be condensed into a single protein.

## Introduction

Central to our understanding of immunity and host defense are the signaling pathways of the innate immune system. Prototypical examples include the pathways activated by the Toll-like Receptors (TLRs), RIG-I like Receptors (RLRs) and the Nucleotide binding Leucine Rich Repeat containing (NLR) proteins. Upon detection of microbial products, virulence factors or dysregulation of cellular homeostasis, members of these diverse receptor families seed the assembly of multiprotein complexes known as supramolecular organizing centers (SMOCs; (Kagan et al., 2014). SMOCs represent the signaling organelles of the innate immune system, which unleash biochemical activities that promote inflammation, interferon responses, changes in metabolism or cell death, in a context-dependent manner. Examples of these signaling organelles include the inflammasomes, which serve as the subcellular sites of interleukin-1β (IL-1β) maturation and signals that induce pyroptosis (Schroder and Tschopp, 2010).

Inflammasomes are controlled by caspases that operate upstream or downstream of these molecular machines, including caspase-1 (Casp-1), murine Casp-11 (mCasp-11) or human Casp-4 (hCasp-4) and −5 (Kesavardhana et al., 2020). These enzymes share a common domain architecture, consisting of an N-terminal caspase activation and recruitment domain (CARD) fused to an enzymatic domain. Upon activation, inflammatory caspases cleave the cytosolic protein gasdermin D (GSDMD), which subsequently forms pores on the plasma membrane to cause lytic cell death (pyroptosis) and the release of the cleaved IL-1 family cytokines IL-1β and IL-18 (Kayagaki et al., 2015; Liu et al., 2016; Shi et al., 2015).

Despite their commonalities, inflammatory caspases possess protein-specific activities. For example, only Casp-1 has considerable IL-1 converting enzyme (ICE) activity (Ramirez et al., 2018) and only Casp-1 can be recruited into inflammasomes in order to stimulate its catalytic activity (Schroder and Tschopp, 2010). mCasp-11, hCasp-4 and −5, in contrast, are not recruited into inflammasomes and their catalytic activity is stimulated by binding of their CARD to a microbial product—bacterial lipopolysaccharide (LPS) (Shi et al., 2014). As mCasp-11 and hCasp-4 cannot cleave pro-IL-1β, the pathways activated by LPS depend on the downstream activation of the NLRP3 inflammasome. Inflammasome-associated Casp-1 then provides the ICE activity that the upstream caspases cannot (Baker et al., 2015; Rühl and Broz, 2015). Notably, no caspase is known to combine LPS-binding and IL-1β processing activities. Despite the importance of each of the inflammatory caspases in host defense, the mechanisms underlying their differential cleavage specificities and ligand-binding activities are poorly defined.

Core components of the pyroptosis machinery are conserved throughout vertebrate evolution. Mammals of the order *Carnivora*, including all terrestrial and marine dog-like and cat-like animals, represent an exception to this statement. These animals lack the gene encoding Casp-1 (Eckhart et al., 2008, 2009). Instead, carnivorans possess a gene where a Casp-1-like CARD is fused to a second CARD and an enzymatic domain, both of which are most similar to hCasp-4 (Figure 1A). In addition, transcripts of this gene are alternatively spliced to give rise to two isoforms: one that contains a single CARD (Casp-1/4a) and one with both CARDs (Casp-1/4b) (Figure 1A). Importantly, both carnivoran caspases contain enzymatic domains that are most similar to hCasp-4, which does not contain ICE activity. These bioinformatic observations raise the question of how pyroptosis and IL-1 release is regulated in carnivorans, as key elements of the known pathways are reorganized or missing.

**Figure 1:**
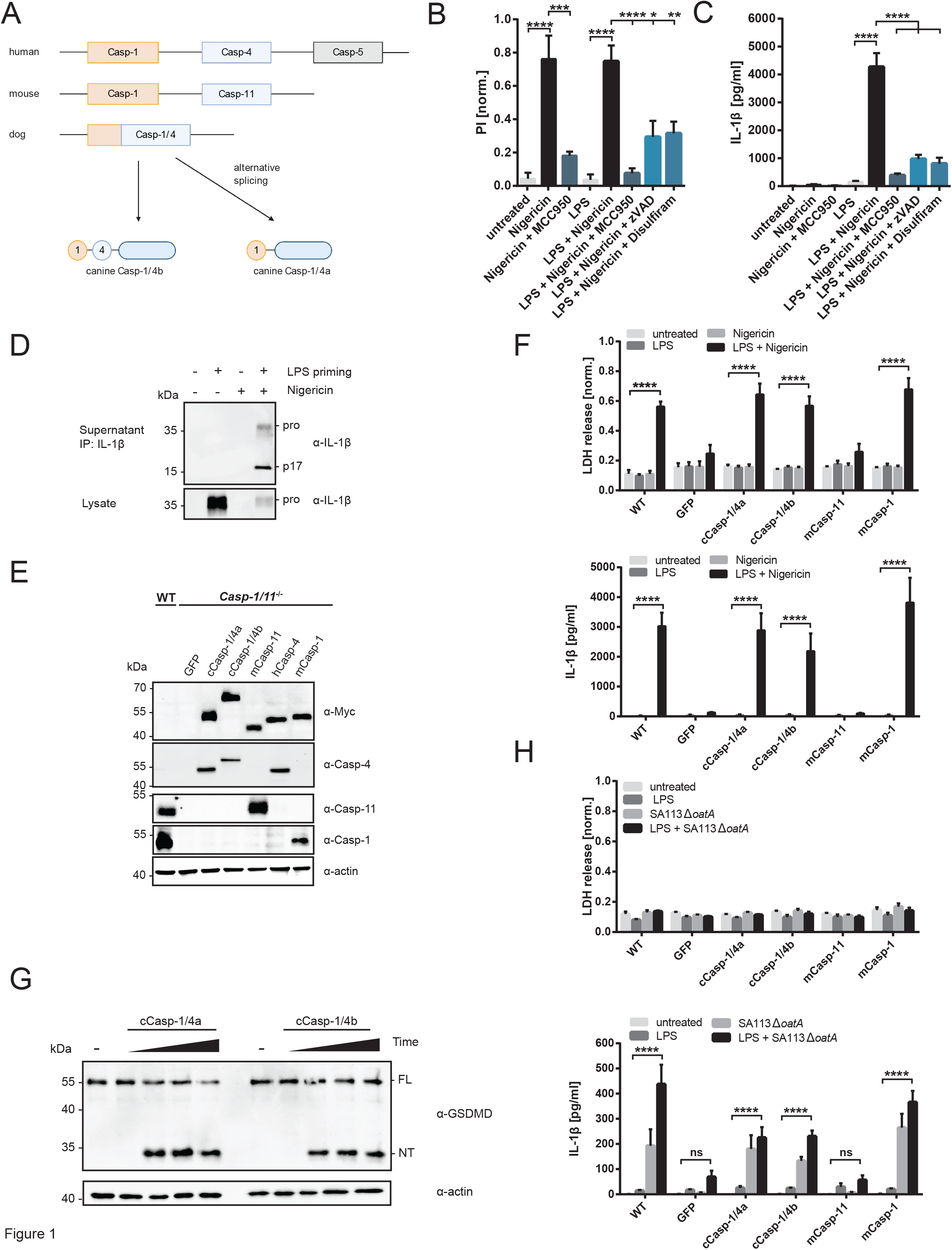
Canine Casp-1/4 proteins can functionally complement for Casp-1 in the context of inflammasome activation. (A) Schematic representation of genetic loci encoding inflammatory caspases in humans, mice and dogs. (B-D) Canine primary MDMs were primed with 1 μg/ml of LPS for 4 h or left unprimed before stimulation with Nigericin (10 μM) for 3h. Cells were pre-treated with indicated inhibitors for 30 min and inhibitors were co-administered during nigericin treatment. (B) PI fluorescence intensity as a measure of plasma membrane rupture or pore-formation was quantified after 3 h of nigericin stimulation. (C) Quantification of released IL-1β from cell-free supernatants was performed by ELISA after 3 h. (D) Immunoblot analysis of immunoprecipitated IL-1β from cell-free supernatants and cell-associated IL-1β after 3 h of nigericin treatment. (E) Immunoblot analysis of whole-cell lysates of WT iBMDMs (after priming with 1 μg/ml of LPS for 4h) and *Casp-1/11^−/−^* iBMDMs reconstituted with the indicated caspase. All constructs carry an N-terminal Myc-tag for detection. (F) WT iBMDMs or *Casp-1/11^−/−^* iBMDMs reconstituted with the indicated caspase were primed for 4 h with LPS (1 μg/ml) or left unprimed before treatment with nigericin (10 μM) for 3 h. LDH and IL-1β in cell culture supernatants were then quantified using a colorimetric assay and ELISA, respectively. (G) *Casp-1/11^−/−^* iBMDMs reconstituted with canine casp-1/4 hybrids were primed for 4 h with LPS (1 μg/ml), then either left untreated or stimulated with nigericin (10 μM) for 0 min, 1 h, 2 h, or 3 h. Reactions were stopped by adding concentrated SDS loading buffer directly into the cell culture well to capture proteins in both the cell lysates and the cell culture supernatant. Proteins were separated by SDS-PAGE and processing of GSDMD was analyzed by immunoblotting. (H) WT iBMDMs or *Casp-1/11^−/−^* iBMDMs reconstituted with the indicated caspase were primed for 4 h with LPS (1 μg/ml) or left unprimed before infection with *S. aureus* 113 Δ*oatA* at an MOI of 30. LDH and IL-1β in cell culture supernatants were quantified 12 h post infection using a colorimetric assay and ELISA, respectively. Data are represented as mean ± SEM of at least three independent experiments. Immunoblots show representative result of three independent repeats. Statistical significance was determined by one-way ANOVA (B+C) or two-way ANOVA (F+H): *p < 0.05; **p < 0.01; ***p < 0.001; ****p < 0.0001.

Herein, we present a detailed characterization of the Casp-1/4 proteins from the common dog (*Canis lupus familiaris*). We show that these enzymes, despite their evolutionary origin as Casp-4 homologues, display all activities of Casp-1, including the ability to cleave pro-IL-1β. Comparative analysis revealed how human and murine caspase-1 selects pro-IL-1β as a substrate, by a mechanism distinct from that which detects GSDMD. This knowledge enabled us to redesign hCasp-4 into a protease exhibiting substantial ICE activity *in vitro* and in cells. Within macrophages, redesigned hCasp-4 operated as a one-protein signaling pathway that directly binds LPS and cleaves IL-1 and GSDMD, by a process that bypasses the need for inflammasomes. Remarkably, a broader evolutionary analysis revealed that cats encode a Casp-4 gene whose product naturally operates in a similar manner to our redesigned hCasp-4. These findings reveal molecular determinants of caspase substrate specificity and challenge the idea that complexity is a prerequisite for innate immune pathway design.

## Results

### Despite bioinformatic predictions, canine inflammatory caspases are functional homologues of Casp-1

The gene encoding the inflammasome stimulatory protein NLRP3, which responds to disturbances of cellular homoeostasis indicated by ROS production, lysosomal damage or aberrant ion fluxes, is conserved in carnivorans (Hornung et al., 2008; Muñoz-Planillo et al., 2013; Perregaux and Gabel, 1994; Pétrilli et al., 2007; Zhou et al., 2011). NLRP3-induced pyroptosis is usually described as a two-step process: After transcriptional priming with a TLR-ligand (signal 1), which leads to the upregulation of NLRP3 and pro-IL-1β, a secondary signal drives NLRP3-mediated inflammasome activity and IL-1β maturation and release (signal 2).

To determine if NLRP3 is operational in carnivoran cells, we isolated primary canine monocytes from whole blood and generated monocyte-derived macrophages (MDMs) (Figure S1A). These cells were primed with LPS and stimulated with the K^+^ ionophore nigericin, an inducer of NLRP3 in murine and human cells. Stimulated cells were then stained with the membrane impermeable dye propidium iodide (PI), which binds to intracellular nucleic acids upon plasma membrane disruption, and determined cellular ATP levels as a proxy for viability. Treatments of canine MDMs with nigericin stimulated an increase in PI fluorescence and a decrease in cellular ATP, both of which are indicators of cell death (Figure 1B; Figure S1B). Nigericin induced these responses in the presence or absence of LPS priming, a finding that could be explained by constitutive expression of NLRP3, as is observed in human primary monocytes (Gritsenko et al., 2020). When cells were primed with LPS, these responses coincided with the release of IL-1β into the cell culture supernatant (Figure 1C). Immunoprecipitation and immunoblot analysis of extracellular IL-1β confirmed that this cytokine was processed into the bioactive p17 fragment (Figure 1D; fragment at ~17 kDa). The NLRP3 inhibitor MCC950 (Coll et al., 2015) prevented nigericin-induced cell death, as cells treated with this inhibitor exhibited lower PI staining and higher ATP levels, as compared to non-inhibitor-treated cells (Figure 1B; Figure S1B). In addition, the pan-caspase inhibitor zVAD-FMK and disulfiram, a specific inhibitor of GSDMD pore formation (Hu et al., 2020), led to a reduction of nigericin-induced death in canine MDMs. (Figure 1B; Figure S1B). All three inhibitors diminished IL-1β release from LPS-primed cells when co-administered during nigericin stimulation (Figure 1C). Overall, these data support the idea that the NLRP3 inflammasome pathway is intact in primary canine cells.

Our finding that IL-1β can be cleaved and released from canine MDMs was notable, as Casp-1 is missing in dogs and other carnivorans. To explain these findings, we determined if canine Casp-1/4 (cCasp-1/4a and cCasp-1/4b) proteins can functionally operate as Casp-1. We designed an experimental system based on stable, heterologous expression of a caspase in immortalized bone marrow-derived macrophages (iBMDMs) from mice deficient in mCasp-1 and mCasp-11 (hereafter referred to as *Casp-1/11^−/−^* iBMDMs). We reconstituted *Casp-1/11^−/−^* iBMDMs with a panel of inflammatory caspases from different species, including both cCasp-1/4 isoforms (cCasp-1/4a and cCasp-1/4b), murine Casp-1 (mCasp-1) and Casp-11 (mCasp-11), and human Casp-4 (hCasp-4). Notably, cCasp-1/4a and cCasp-1/4b could be detected by immunoblot using a Casp-4-specific, but not a Casp-1-specific, antibody, underscoring their identity as structural Casp-4 homologues (Figure 1E). As the parental iBMDMs lack mCasp-1 and mCasp-11, any observed pyroptotic phenotypes upon treatment with inflammasome triggers are attributed to caspases introduced as transgenes.

We primed cells with LPS and subsequently stimulated with nigericin. Within wild type (WT) cells, these treatments induced lytic cell death, as assessed by the release of the cytosolic enzyme lactate dehydrogenase (LDH) into the cell culture supernatant, and secretion of IL-1β (Figure 1F). Both processes were abrogated in *Casp-1/11^−/−^* iBMDMs expressing GFP or mCasp-11 as a transgene, but were restored by expression of mCasp-1 (Figure 1F). Similar to mCasp-1, cells expressing cCasp-1/4a or cCasp1/4b released LDH and IL-1β upon LPS priming and nigericin treatment. These phenotypes coincided with the cleavage of GSDMD into its pore-forming N-terminal domain (Figure 1G), a hallmark of pyroptotic cell death. Similar results were obtained when cells were transfected with the double-stranded DNA analogue poly(dA:dT) (Figure S1C, D), which stimulates the AIM2 inflammasome (Fernandes-Alnemri et al., 2009; Hornung et al., 2009). Poly(dA:dT)-induced LDH release was independent of LPS priming (Figure S1C), which is consistent with reports that AIM2 is constitutively expressed in murine macrophages (Hornung et al., 2009).

In addition to stimulating pyroptosis, select inflammasome activators can induce IL-1β release from living cells (Zanoni et al., 2016). These cells are known as hyperactive, as they have added IL-1β to the repertoire of cytokines that LPS-activated cells can secrete. Infection of macrophages with *S. aureus* lacking O-acetyltransferase A (SA113 Δ*oatA*) causes hyperactivation in a Casp1 and GSDMD-dependent manner (Evavold et al., 2018; Wolf et al., 2016). To test whether cCasp-1/4 isoforms respond to hyperactivating stimuli, we infected our set of transgene-expressing *Casp-1/11^−/−^* iBMDMs with SA113 Δ*oatA*. mCasp-1-expressing iBMDMs, as well as WT counterparts, responded to infection with the release of IL-1β and the amount of IL-1β could be boosted if cells were primed with LPS (Figure 1H). Cells expressing cCasp-1/4a or cCasp-1/4b responded similarly to infection with the release of IL-1β, although the amount of IL-1β released was lower than with mCasp-1 expressing cells (Figure 1H). As expected, no IL-1β was released from *Casp-1/11^−/−^* iBMDMs expressing GFP or mCasp-11. Also as expected for a stimulus that induces IL-1β release from living cells, no increases in LDH release were observed upon infection of WT or transgene expressing *Casp-1/11^−/−^* iBMDMs (Figure 1H).

Within mCasp-1, the N-terminal CARD is critical for recruitment into inflammasomes via interactions with the upstream adaptor ASC (Srinivasula et al., 2002; Stehlik et al., 2003). To determine the role of the N-terminal CARD present in cCARD-1/4a and cCARD1/4b, we reconstituted *Casp-1/11^−/−^* iBMDMs with a canine caspase construct lacking its Casp-1-like CARD. Cells expressing this mutant caspase (cCasp-1/4bΔCARD) failed to release LDH or IL-1β after LPS priming followed by nigericin treatment (Figure S1E-H). These collective data demonstrate that both cCasp-1/4 isoforms can operate similar to Casp-1 in the context of multiple inflammasome stimuli, and that their N-terminal CARD is critical for this activity.

### Canine caspases are not LPS sensors and canine cells cannot respond to intracellular LPS

cCasp-1/4b is the only inflammatory caspase known that contains two N-terminal CARDs. While the first CARD is similar to that of Casp-1, the second CARD shares homology with the LPS-binding CARDs of hCasp-4 and mCasp-11. We therefore investigated whether cCasp-1/4b might respond to exposure to LPS. Transgenic cells were primed with extracellular LPS before delivery of LPS into the cytosol via electroporation. As expected, LPS electroporation stimulated LDH release from WT iBMDMs and *Casp-1/11^−/−^* iBMDMs expressing mCasp-11 or hCasp-4, indicating the induction of pyroptosis (Figure 2A). Cells expressing mCasp-1 or GFP did not lyse upon LPS electroporation, as neither of these proteins can bind LPS. Notably, none of our reconstituted *Casp-1/11^−/−^* iBMDMs released significant amounts of IL-1β after LPS electroporation (Figure 2B). This finding validates current dogma, which predicts that mCasp-11 and mCasp-1 are both needed for LPS to induce cleavage and release IL-1β (Rühl and Broz, 2015). Since our iBMDMs produce either mCasp-1 or mCasp-11 (not both), IL-1β release is diminished.

**Figure 2:**
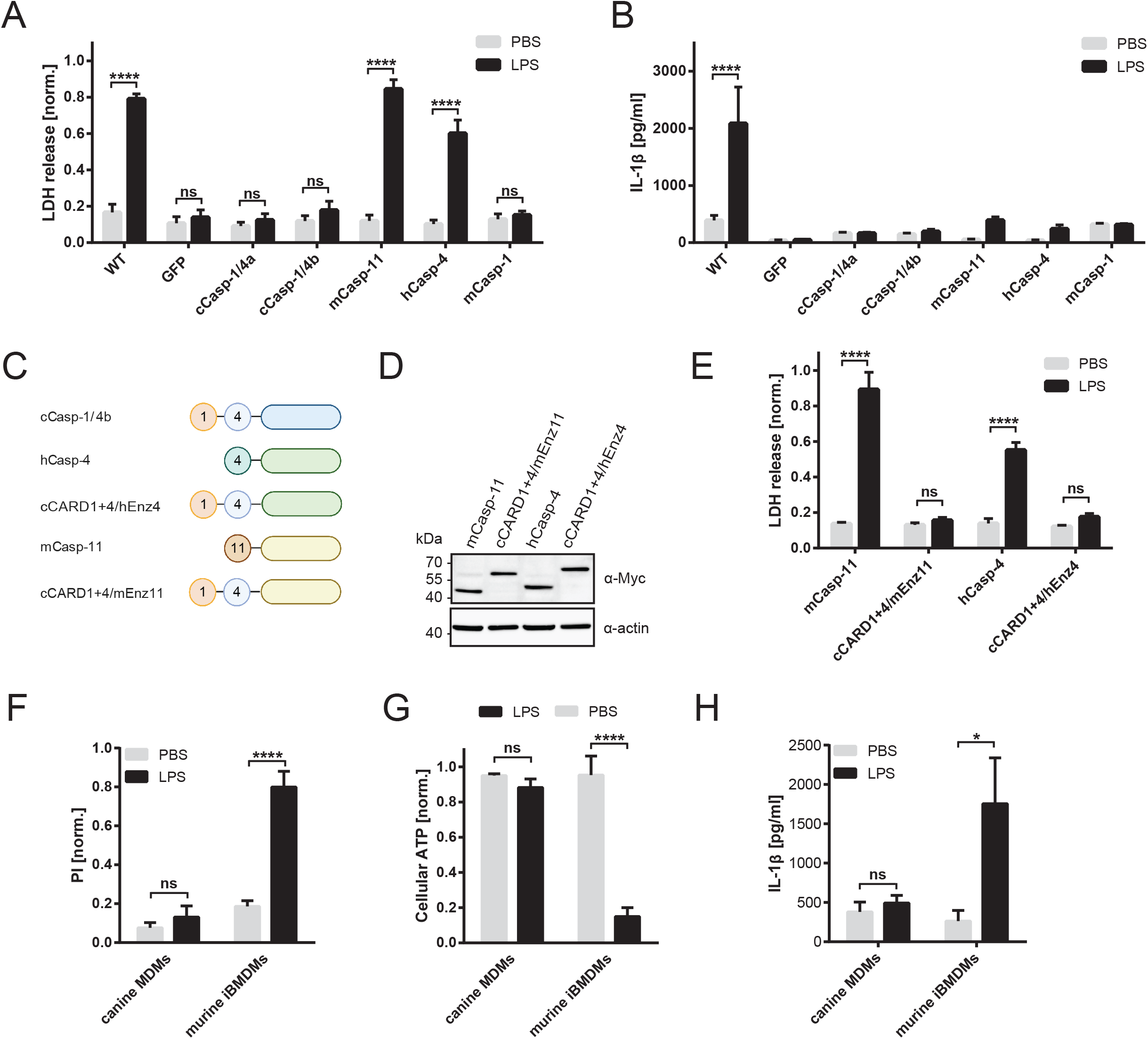
Canine Casp-1/4 proteins do not act as cytosolic LPS sensors. (A) WT iBMDMs or *Casp-1/11^−/−^* iBMDMs reconstituted with the indicated caspase were primed for 4 h with LPS (1 μg/ml) before electroporation of LPS or PBS as negative control. LDH in cell-free supernatants was quantified after 3 h using a colorimetric assay. (B) WT iBMDMs or *Casp-1/11^−/−^* iBMDMs reconstituted with the indicated caspase were primed for 4 h with LPS (1 μg/ml) before electroporation of LPS or PBS as negative control. Levels of IL-1β in cell-free supernatants were quantified after 3 h by ELISA. (C) Schematic representation of WT and chimeric caspase constructs. (D) Immunoblot analysis of whole-cell lysates of *Casp-1/11^−/−^* iBMDMs reconstituted with the indicated WT or chimeric caspase to confirm equal expression levels of all constructs. All constructs carry an N-terminal Myc-tag for detection. (E) *Casp-1/11^−/−^* iBMDMs reconstituted with the indicated WT or chimeric caspase were primed for 4 h with LPS (1 μg/ml) or left unprimed before electroporation of LPS or PBS as negative control. LDH in cell-free supernatants was quantified after 3 h using a colorimetric assay. (F-H) Canine primary MDMs or murine WT iBMDMs were primed with 1 μg/ml of LPS for 4 h. Subsequently, 1 μg of LPS was delivered into the cytosol of the cells by electroporation. PBS electroporation served as negative controls. (F) PI fluorescence intensity as a measure of plasma membrane rupture or pore-formation was quantified 3 h post-electroporation. (G) Intracellular ATP levels were determined after 3 h using Celltiter-Glo viability assay. (H) IL-1β levels in the extracellular media was quantified by ELISA at 3 h post-electroporation. Data are represented as mean ± SEM of three (E), four (F-H), or five (A+B) independent experiments. Statistical significance was determined by two-way ANOVA: *p < 0.05; **p < 0.01; ***p < 0.001; ****p < 0.0001.

We found that neither cCasp-1/4a nor cCasp-1/4b expressing cells died upon cytosolic LPS delivery (Figure 2A). This finding was surprising, since cCasp-1/4b contains a CARD that is similar to the LPS-binding CARD from hCasp-4. Since cCasp-1/4a and cCasp-1/4b are able to cleave murine GSDMD (Figure 1G), these results imply that cCasp-1/4b cannot become activated by LPS. To further investigate this lack in LPS responsiveness, we determined if the Casp-4-like CARD from cCasp-1/4b can functionally replace the LPS-binding CARD from either mCasp-11 or hCasp-4. To this end, we designed chimeric caspases where the CARDs from cCasp-1/4b were attached to the enzymatic domain from mCasp-11 (mEnz11) or hCasp-4 (hEnz4) (cCARD1+4/mEnz11 and cCARD1+4/hEnz4). These chimeric proteins were stably produced in *Casp-1/11^−/−^* iBMDMs (Figure 2C, D). Cells expressing these chimeras failed to promote pyroptosis when electroporated with LPS (Figure 2E). We further considered the possibility that the Casp-1-like CARD, which is upstream of the Casp-4 like CARD, would prevent LPS detection by cCasp-1/4b. However, deletion of this additional CARD does not render cells responsive to LPS electroporation (Figure S2A). These findings suggest that the Casp-4-like CARD found in cCasp-1/4b is unable to induce pyroptosis in response to LPS. Even within primary canine MDMs, we observed no evidence of cell death or IL-1β release upon LPS electroporation (Figure 2F-H). Altering electroporation conditions did not reveal any LPS-specific change in canine MDMs (Figure S2B-D). In human and pig monocytes (but not MDMs), the TLR4 pathway activates the NLRP3 inflammasome to promote IL-1β release (Gaidt et al., 2016). We observed similar responses when we stimulated canine monocytes with LPS, as these treatments promoted IL-1β release (Figure S2E, F). Canine cells are therefore not generally unresponsive to LPS but are rather specifically unresponsive to cytosolic LPS. This latter observation is likely explained by our finding that cCasp-1/4b cannot respond to LPS within canine cells or upon heterologous expression in murine cells.

### Evolutionary symmetry between the mechanisms of Casp-1-like enzyme activity

Our findings suggest that cCasp-1/4 proteins operate most similarly to Casp-1 homologues in humans and mice. We therefore investigated mechanisms that underlie this symmetry of activities. Cysteine 285 (C285) is the main catalytic residue within human and murine Casp-1 (Wilson et al., 1994), which is conserved in the canine inflammatory caspases. We introduced inactivating Cys-to-Ala mutations at the equivalent sites in cCasp-1/4a and cCasp-1/4b (C285A and C370A, respectively) and expressed these mutants in *Casp-1/11^−/−^* iBMDMs (Figure 3A). Notably, we did not observe any LDH or IL-1β release after LPS + nigericin treatment of cells expressing these cCasp-1/4 mutants (Figure 3B, C). Accordingly, no processed IL-1β was released from these cells (Figure 3D, E), and we did not observe any processing of GSDMD (Figure 3F). These results indicate that the catalytic activity of cCasp-1/4a and cCasp-1/4b is required to induce pyroptosis and IL-1β release.

**Figure 3:**
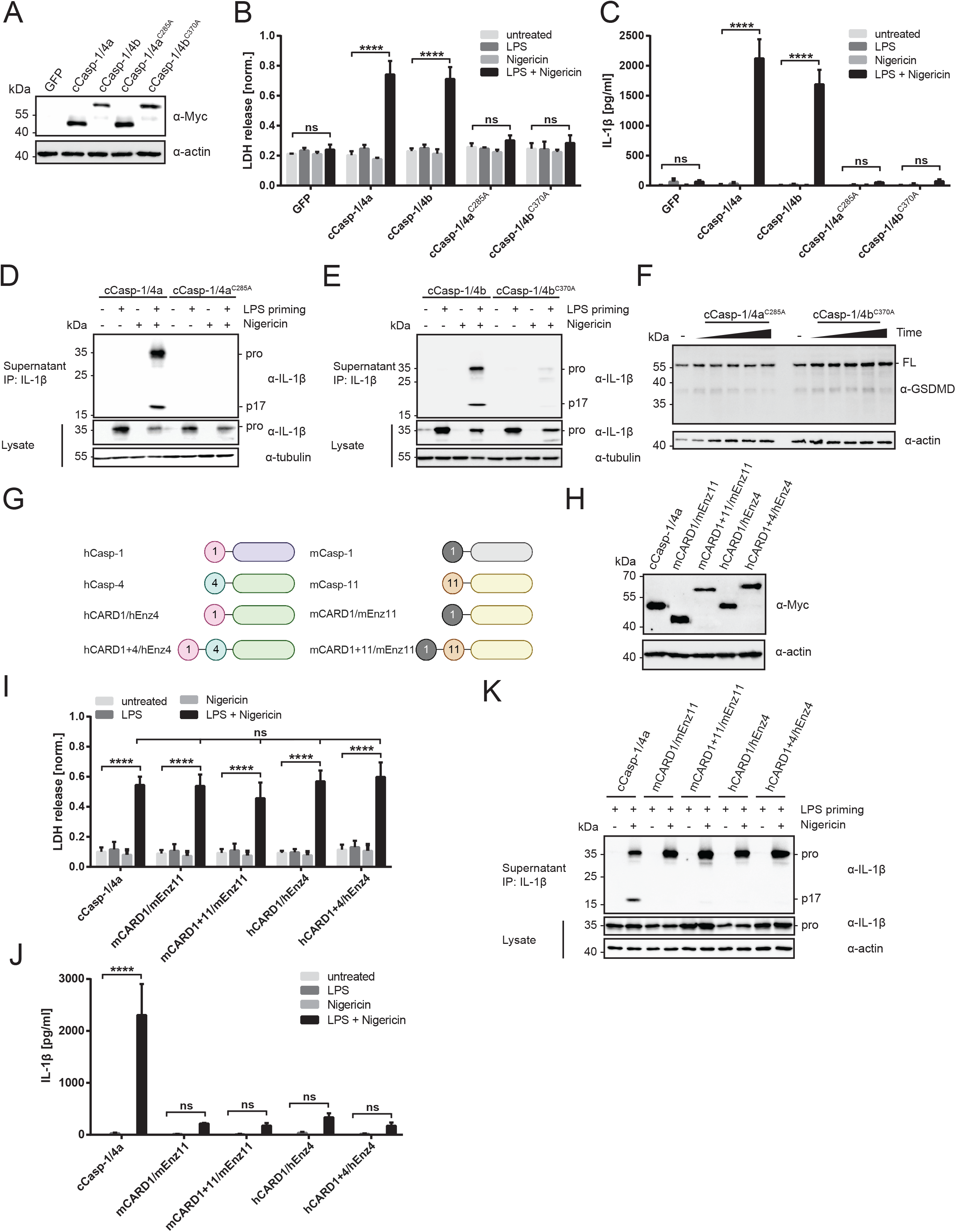
The catalytic activity of cCasp-1/4 is required for inflammasome-dependent IL-1β release. (A) Immunoblot analysis of whole-cell lysates of *Casp-1/11^−/−^* iBMDMs expressing either WT cCasp-1/4 isoforms or catalytically dead cCasp-1/4 variants. All constructs carry an N-terminal Myc-tag for detection. (B, C) *Casp-1/11^−/−^* iBMDMs reconstituted with the indicated caspase were primed for 4 h with LPS (1 μg/ml) or left unprimed before treatment with nigericin (10 μM) for 3 h. LDH and IL-1β in cell culture supernatants were then quantified using a colorimetric assay and ELISA, respectively. (D, E) *Casp-1/11^−/−^* iBMDMs reconstituted with the indicated caspase were primed for 4 h with LPS (1 μg/ml) or left unprimed before stimulation with nigericin (10 μM). Immunoprecipitated IL-1β from cell-free supernatants and cell-associated pro-IL-1β in whole-cell lysates were analyzed by immunoblot after 3 h of nigericin treatment. (F) *Casp-1/11^−/−^* iBMDMs reconstituted with catalytic cCasp-1/4 mutants were primed for 4 h with LPS (1 μg/ml), then either left untreated or stimulated with nigericin (10 μM) for 0 min, 30 min, 1 h, 2 h, or 3 h. Reactions were stopped by adding concentrated SDS loading buffer directly into the cell culture well to capture proteins in both the cell lysates and the cell culture supernatant. Proteins were separated by SDS-PAGE and processing of GSDMD was analyzed by immunoblotting. (G) Schematics showing architecture of synthetic hybrid caspases consisting of human or murine Casp-1 CARDs and the catalytic domains and CARDs of hCasp-4 or mCasp-11, respectively. (H) Immunoblot analysis of whole-cell lysates of *Casp-1/11^−/−^* iBMDMs expressing indicated natural or synthetic hybrid caspases. All constructs carry an N-terminal Myc-tag for detection. (I+J) *Casp-1/11^−/−^* iBMDMs reconstituted with the indicated hybrid caspase were primed for 4 h with LPS (1 μg/ml) or left unprimed before treatment with nigericin (10 μM) for 3 h. LDH and IL-1β in cell culture supernatants were then quantified using a colorimetric assay and ELISA, respectively. (K) *Casp-1/11^−/−^* iBMDMs reconstituted with the indicated hybrid caspase were primed for 4 h with LPS (1 μg/ml) before stimulation with nigericin (10 μM). Immunoprecipitated IL-1β from cell-free supernatants and cell-associated pro-IL-1β in whole-cell lysates were analyzed by immunoblot after 3 h of nigericin treatment. Data are represented as mean ± SEM of three independent experiments. Immunoblots of immunoprecipitated IL-1β display one representative result of three independent repeats. Statistical significance was determined by two-way ANOVA: ****p < 0.0001.

The ability of cCasp-1/4a and cCasp-1/4b to induce the processing of IL-1β was surprising, as their catalytic domain is most similar to that found in hCasp-4, a caspase that has no considerable ICE activity (Bibo-Verdugo et al., 2020; Faucheu et al., 1995). The presence of cleaved IL-1β in the supernatants of the reconstituted macrophages could be explained if all Casp-4 homologues actually do have ICE activity, but this activity can only be stimulated upon recruitment into an inflammasome.

To address this possibility, we created a scenario whereby the enzymatic domains of hCasp-4 or mCasp-11 can be recruited into an inflammasome via a mechanism similar to that which recruits Casp-1. This was accomplished by generating fusions between the CARD of hCasp-1 or mCasp-1 to either full-length hCasp-4 or mCasp-11, or the isolated enzymatic domains of these caspases (Figure 3G). When expressed in *Casp-1/11^−/−^* iBMDMs, these chimeric enzymes caused a similar degree of cell lysis after LPS + nigericin treatment, indicating productive interactions with the inflammasome machinery (Figure 3H, I). However, this pyroptotic cell death was not accompanied by the release of mature IL-1β (Figure 3J, K). These findings eliminate the possibility that recruitment of hCasp-4 or mCasp-11 into an inflammasome stimulates a latent ICE activity.

### Canine caspase activities reveal how IL-1β is selected as a substrate by mCasp-1

Based on the above-described phenotypes, we considered the possibility that cCasp-1/4 enzymes display intrinsic ICE properties. We therefore expressed and purified catalytic domains from mCasp-1, hCasp-4 and cCasp-1/4 from *E. coli.* Analysis by SDS-PAGE confirmed that purified enzymes are autocatalytically processed into the large (p20) and small (p10) catalytic subunits that are important for catalytic activity (Figure 4A). We then characterized their ability to cleave peptide-based and full-length protein substrates *in vitro*. Caspase cleavage sites within substrates have been defined by tetrapeptides, consisting of the four amino acids C-terminal of the scissile peptide bond (Julien and Wells, 2017). Enzyme kinetic analyses using YVAD-pNA, a chromogenic tetrapeptide substrate optimized for cleavage by Casp-1, revealed similar Michaelis-Menten constants (K_m_) for mCasp-1 and cCasp-1/4, while the K_m_ of hCasp-4 for this substrate was so high it was not possible to calculate (Figure 4B, C). The turnover number (k_cat_) and catalytic efficiency of mCasp-1 were only 2-3-fold higher than those of cCasp-1/4 (Figure 4C). This result suggests a shift of cCasp-1/4 from a Casp-4 to a Casp-1-like specificity. This possibility was confirmed when we examined the ability of the caspases to process pro-IL-1β *in vitro*. A serial dilution of the enzyme of interest was incubated with a fixed amount of murine pro-IL-1β and substrate cleavage was quantified by immunoblot to obtain catalytic efficiencies. As expected, pro-IL-1β was cleaved far more efficiently by mCasp-1 than by hCasp-4 (Figure 4D, E). Intriguingly, cCasp-1/4 was only slightly less efficient at cleaving pro-IL-1β than mCasp-1 (Figure 4F). Quantification revealed that hCasp-4 was more than 600 times less efficient at cleaving pro-IL-1β than mCasp-1 and cCasp-1/4 (Figure 4G). This finding demonstrates that cCasp-1/4 enzymes have the ability to process pro-IL-1β at a near-comparable rate to that of mCasp-1.

**Figure 4:**
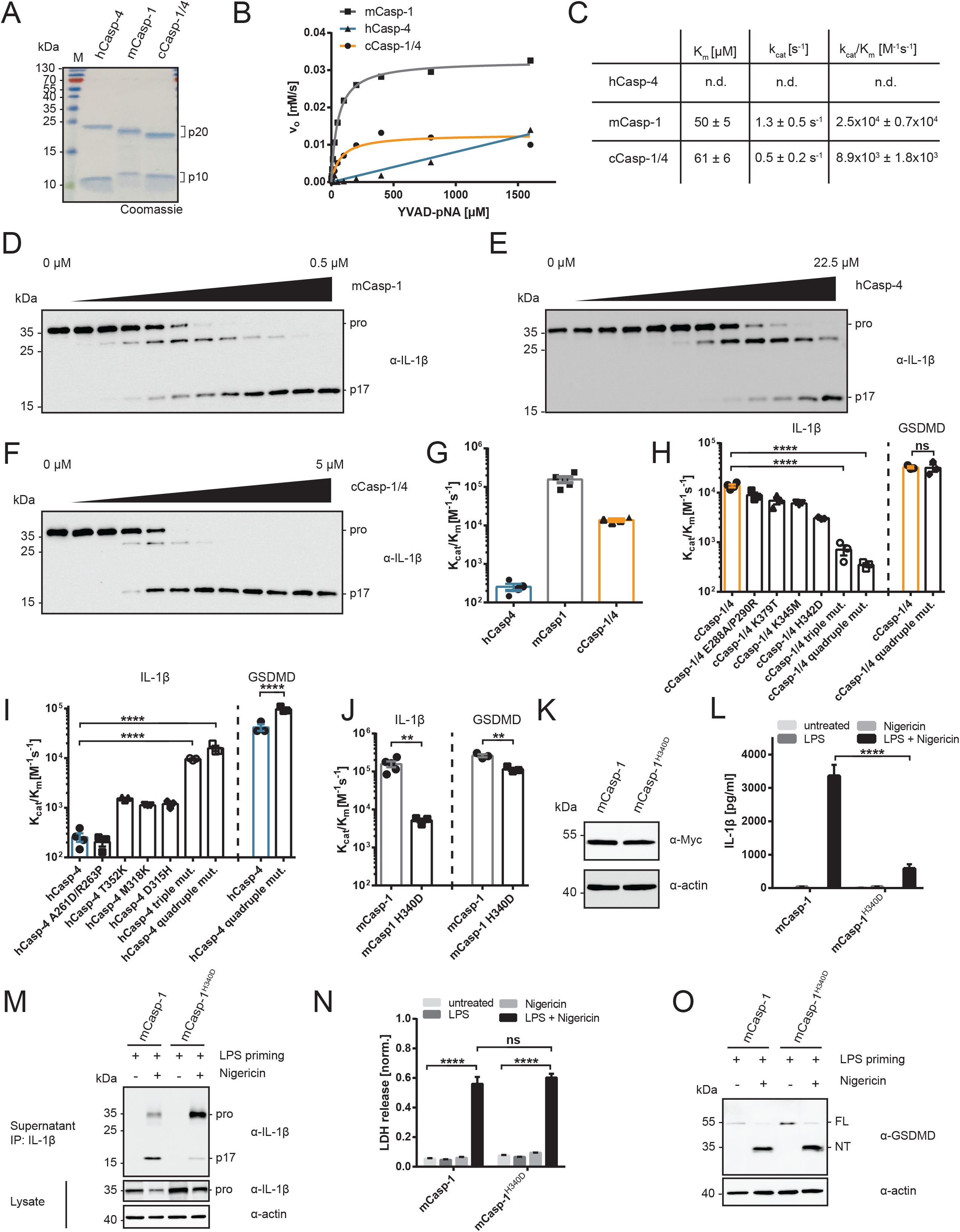
Mechanistic insights in how inflammatory caspases select IL-1β as a substrate. (A)Catalytic domains of hCasp-4, mCasp-1 and cCasp-1/4 were expressed in *E. coli.* Purified proteins were separated by SDS-PAGE and visualized by Coomassie staining to confirm autocatalytic processing and purity. (B, C) Enzyme kinetics analysis of the cleavage of the chromogenic Casp-1-substrate YVAD-pNA by recombinant purified caspases. (B) Purified caspases (20 nM) were mixed with serially diluted substrate (0 – 1600 μM), initial velocities were plotted in dependence of the substrate concentration and fitted according to the hyperbolic Michaelis-Menten equation to derive kinetic parameters. (C) Calculated kinetic parameters of YVAD-pNA cleavage reaction for each tested caspase. Parameters for hCasp-4 could not be accurately determined as curves did not reach saturation in the tested range of substrate concentrations. Displayed parameters represent mean ± SD of three independent experiments involving two separate caspase preparations. (D-G) 50 nM of recombinant murine pro-IL-1β was mixed with two-fold serial dilutions of purified caspase catalytic domains and incubated for 30 min at 37 °C. Proteins were then separated by SDS-PAGE and processing of pro-IL-1β was analyzed by immunoblotting. Highest enzyme concentration used in each assay is indicated above the blot. Catalytic efficiencies (k_cat_/K_m_) were calculated based on disappearance of band corresponding to pro-form of IL-1β. (H) Catalytic efficiencies of *in vitro* pro-IL-1β and GSDMD cleavage by cCasp-1/4 point mutants. Triple mutant carries K379T, K345M and H342D mutations. Quadruple mutant combines these three mutations with E288A/P290R mutation. (I) Catalytic efficiencies of *in vitro* pro-IL-1β and GSDMD cleavage by hCasp-4 point mutants. Triple mutant carries T352K, M318K and D315H mutations. Quadruple mutant combines these three mutations with A261E/R263P mutation. (J) Catalytic efficiencies of *in vitro* pro-IL-1β and GSDMD cleavage by mCasp-1 H340D point mutant compared to WT mCasp-1. (K) Immunoblot analysis of whole-cell lysates of *Casp-1/11^−/−^* iBMDMs expressing either WT mCasp-1 or mCasp-1 with a H340D mutation. Both constructs are N-terminally Myc-tagged to enable detection. (L, N) *Casp-1/11^−/−^* iBMDMs reconstituted with mCasp-1 or mCasp-1 H340D were primed for 4 h with LPS (1 μg/ml) or left unprimed before treatment with nigericin (10 μM) for 3 h. LDH and IL-1β in cell culture supernatants were then quantified using a colorimetric assay and ELISA, respectively. (M) *Casp-1/11^−/−^* iBMDMs reconstituted with WT mCasp-1 or mCasp-1 H340D were primed for 4 h with LPS (1 μg/ml) or left unprimed before stimulation with nigericin (10 μM). Immunoprecipitated IL-1β from cell-free supernatants and cell-associated pro-IL-1β in whole-cell lysates were analyzed by immunoblotting after 3 h. (O) *Casp-1/11^−/−^* iBMDMs reconstituted with mCasp-1 or mCasp-1 H340D were primed for 4 h with LPS (1 μg/ml), then either left untreated or stimulated with nigericin (10 μM) for 3 h. Reactions were stopped by adding concentrated SDS loading buffer directly into the cell culture well to capture proteins in both the cell lysates and the cell culture supernatant. Proteins were separated by SDS-PAGE and processing of GSDMD was analyzed by immunoblotting. Each data point (G-J) represents the result of one independent assay. Bars and error bars represent mean ± SEM of at least three independent experiments. Repeats of *in vitro* assays involved at least two independent caspase preparations. Immunoblots are representative of at least three independent repeats. Statistical significance was determined by one-way ANOVA (H-J) or two-way ANOVA (L+N): *p < 0.05; **p < 0.01; ***p < 0.001; ****p < 0.0001.

To identify the molecular determinants for the ICE activity of cCasp-1/4 enzymes, we first generated chimeric catalytic domains consisting of the p20 domain of cCasp-1/4 and the p10 domain of hCasp-4, and vice versa. Both chimeric catalytic domains exhibited intermediate catalytic efficiencies of pro-IL-1β cleavage (Figure S3A-C). This finding suggests that sites distributed across both the large and small subunit contribute to pro-IL-1β cleavage, with sites in the small subunit being more dominant.

We reasoned that critical residues should be conserved among carnivoran Casp-1/4 homologues, but not in hCasp-4. By applying this rationale, we identified several highly conserved residues (E288/P290, H342, K345, K379) in cCasp-1/4, which we individually replaced with the respective amino acids from hCasp-4. Each of the single point mutations (E288A/P290R, H342D, K345M, K379T) led to a slight decrease in cleavage efficiency compared to WT cCasp-1/4 (Figure 4H). However, when combining three (triple mutant: H342D, K345M, and K379T) or all four of the mutations (quadruple mutant), we observed additive effects, causing pro-IL-1β cleavage efficiency to drop to a level similar to hCasp-4 (Figure 4H; Figure S3E-J). Notably, the ability of the quadruple mutant to cleave GSDMD *in vitro* remained unaffected, indicating a specific impact of the mutations on cleavage of pro-IL-1β, without affecting the overall catalytic activity of the caspase (Figure 4H; Figure S3K, L). Introduction of the reciprocal canine-specific mutations into hCasp-4 had the opposite effect. While individual point mutations led to modest increases (~4-6-fold) in the ability to cleave pro-IL-1β *in vitro*, by combining mutations at several sites we were able to generate a hCasp-4 quadruple mutant. This hCasp-4 quadruple mutant is as efficient as cCasp-1/4 at processing pro-IL-1β (Figure 4I; Figure S4A-H). We observed only minor effects of these mutations on the cleavage of GSDMD *in vitro* (Figure 4I). These findings therefore establish amino acids that determine ICE activity in human and canine caspases.

When mapping the amino acids of interest onto a homology model of the enzymatic domain of cCasp-1/4 in complex with the peptide inhibitor zVAD-FMK, we found that the conserved positively charged residues H342, K345 and K379 are located in the same region in the p10 subunit: a cleft formed by two loops proximal to the tetrapeptide binding site (Figure S4I). This charged site is spatially distinct from the hydrophobic recognition site for GSDMD (highlighted in yellow in Figure S4I) (Liu et al., 2020; Wang et al., 2020). This observation suggests that positive charges in this cavity promote recognition of pro-IL-1β as a substrate for cleavage. To test this model, we created additional caspases carrying charge reversal mutations within this region. A D381R mutation in cCasp-1/4 increased its ability to cleave pro-IL-1β by 4-fold (Figure S5A, B), whereas a R354D mutation in hCasp-4 further decreased its cleavage efficiency by 10-fold (Figure S5A, C). Similar to its canine counterpart, mCasp-1 E379R was ~3-fold better at cleaving pro-IL-1β than WT mCasp-1 (Figure S5A, D).

H342 drew our attention, as it is positioned in a manner that suggests a direct involvement in coordinating the tetrapeptide in the active site (Figure S4I). H342 is present in most carnivoran Casp-1/4 proteins and is strictly conserved in Casp-1. Casp-4 homologues that contain no ICE activity, such as hCasp-4 and mCasp-11, display a negative charge at this site (Figure S5E). Introducing a charge-swap H340D mutation into mCasp-1 greatly decreased its ability to process pro-IL-1β *in vitro*, while only slightly affecting GSDMD cleavage (~31-fold and ~2-fold reduction compared to WT mCasp-1, respectively) (Figure 4J; Figure S5F-H). To determine if this *in vitro* cleavage deficiency extended to activities within cells, we reconstituted *Casp-1/11^−/−^* iBMDMs with a mCasp-1 H340D mutant (Figure 4K). After LPS + nigericin treatment to activate the NLRP3 inflammasome, we detected significantly less IL-1β in the supernatants of cells expressing mCasp-1 H340D, as compared to WT mCasp-1-expressing cells (Figure 4L). Immunoprecipitation of the cytokine from the media confirmed defective processing of IL-1β in mCasp-1 H340D expressing cells (Figure 4M). Importantly, LDH release, GSDMD processing and autocatalytic cleavage of mCasp-1 H340D were unimpaired, indicating an IL-1β-specific effect of the H340D mutation (Figure 4N, O; Figure S5I). These collective findings establish that the charge of amino acids within the catalytic site determines ICE activity, but has minimal impact on the GSDMD-cleaving activity of inflammatory caspases.

### Redesign of hCasp-4 to intrinsically link LPS detection to IL-1β cleavage and release, without the need for inflammasomes

In murine cells, mCasp-11 and mCasp1 are required to link LPS detection to the release of cleaved IL-1β, with inflammasomes serving as an intermediate between these enzymes. Based on the finding that we can redesign hCasp-4 into an enzyme with the ability to process IL-1β *in vitro*, it should be possible to condense the natural multistep LPS-mediated pathway to IL-1β release into one single protein. To fulfill this task, hCasp-4 would need to exhibit LPS-sensing, GSDMD and IL-1β cleavage functionalities. As a proof of concept of this idea, we first designed chimeric caspases consisting of the LPS-binding CARDs from hCasp-4 or mCasp-11 and the enzymatic domain of cCasp-1/4 (named hCARD4/cEnz and mCARD11/cEnz, respectively). These chimeric enzymes were expressed in *Casp-1/11^−/−^* iBMDMs (Figure 5A; Figure S6A), which were then stimulated with extracellular LPS before electroporating them with LPS. Cells expressing either chimeric caspase underwent pyroptosis, as assessed by LDH release after LPS electroporation (Figure 5B, C). Cells expressing either chimeric caspase also released higher amounts of IL-1β, as compared to cells expressing WT caspases (Figure 5D, E). Notably, the cleaved p17 fragment of IL-1β was exclusively detected in the supernatants of cells expressing hCARD4/cEnz or mCARD11/cEnz, but not of cells expressing hCasp-4 or mCasp-11 (Figure 5F, G). The ability of these chimeric caspases to promote IL-1β cleavage and release was independent of NLRP3, as production of this cytokine was insensitive to NLRP3 inhibition by MCC950 (Figure 5H, I; Figure S6B, C). In contrast, MCC950 prevented IL-1β release (but not LDH release) from WT iBMDMs after LPS electroporation (Figure S6D, E), as expected (Baker et al., 2015). These findings demonstrate that caspases can be engineered to bypass the need for inflammasomes to promote IL-1β cleavage and release.

**Figure 5:**
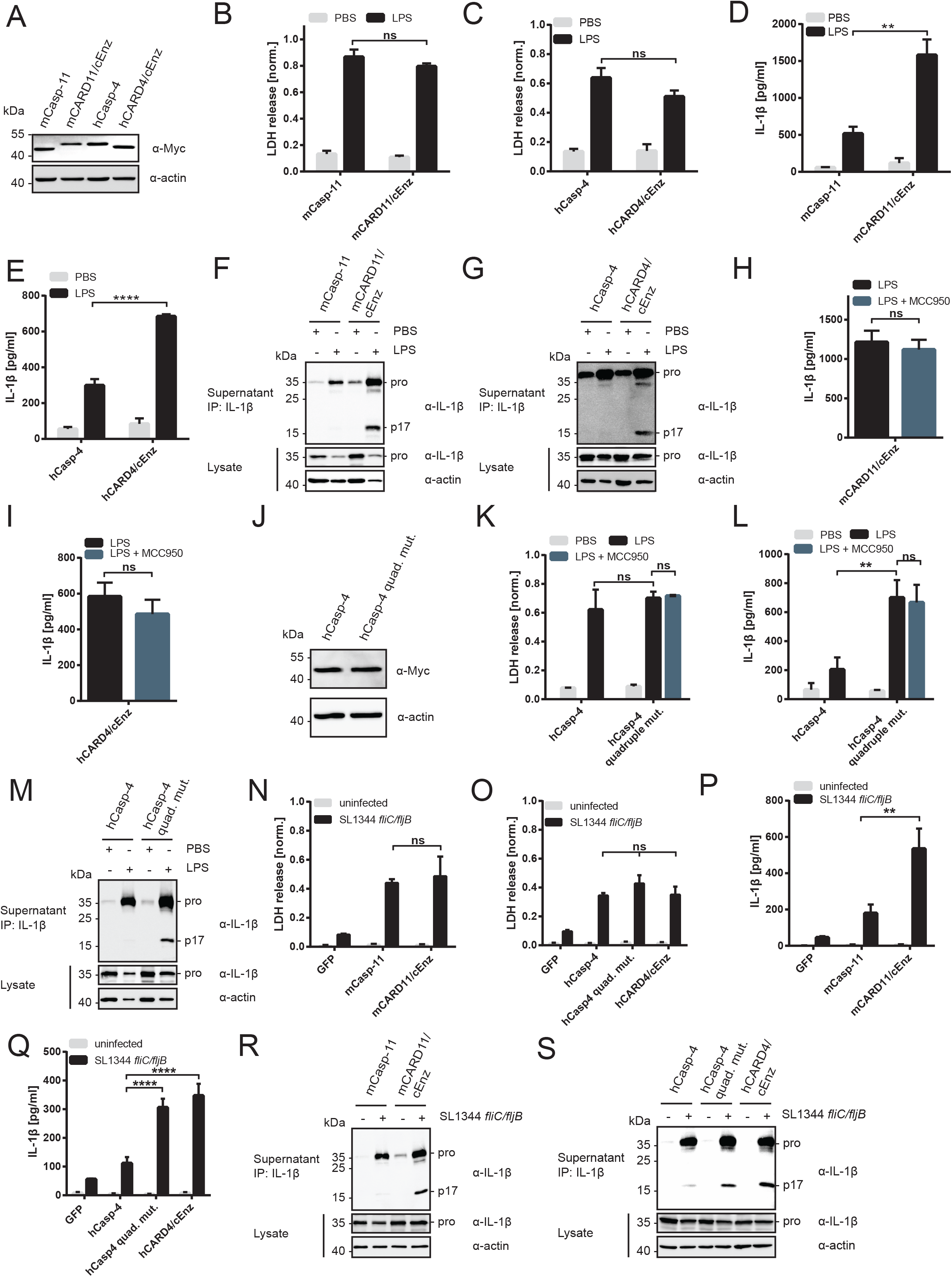
Design of a synthetic one-protein signaling pathway that links LPS detection and IL-1β release, independent of inflammasomes. (A) Immunoblot analysis of whole-cell lysates of *Casp-1/11^−/−^* iBMDMs expressing either mCasp-11, hCasp-4 or chimeric caspases consisting of the CARD domains of hCasp-4 or mCasp-11 and the catalytic domain of cCasp-1/4. All constructs carry an N-terminal Myc-tag for detection. (B,C,D,E,F,H,I,K,L) *Casp-1/11^−/−^* iBMDMs reconstituted with the indicated caspase were primed for 4 h with LPS (1 μg/ml) before electroporation of LPS or PBS as negative control. LDH release and IL-1β levels in cell-free supernatants were quantified after 3 h using a colorimetric assay and ELISA, respectively. In some experiments the NLRP3 inhibitor MCC950 (10 μM) was present in the cell culture media after electroporation. (F,G,M) *Casp-1/11^−/−^* iBMDMs reconstituted with the indicated caspase were primed for 4 h with LPS (1 μg/ml), followed by delivery of LPS (or PBS) into the cytosol via electroporation. Immunoprecipitated IL-1β from cell-free supernatants and cell-associated pro-IL-1β in whole-cell lysates were analyzed by immunoblotting after 3 h. (J) Immunoblot analysis of whole-cell lysates of *Casp-1/11^−/−^* iBMDMs expressing hCasp-4 or hCasp-4 quadruple mutant (T352K/M318K/D315H/ANR261ENP). Both constructs carry an N-terminal Myc-tag for detection. (N-S) *Casp-1/11^−/−^* iBMDMs reconstituted with the indicated caspase were primed for 3 h with LPS (1 μg/ml) before infection with flagellin-deficient Salmonella (SL1344 *fliC/fljB*) at an MOI of 100. LDH release and IL-1β levels in cell-free supernatants were quantified after 4 h using a colorimetric assay and ELISA, respectively. Immunoprecipitated IL-1β from cell-free supernatants and cell-associated pro-IL-1β in whole-cell lysates were analyzed by immunoblotting after 4 h. Data are represented as mean ± SEM of three independent experiments. Immunoblots of immunoprecipitated IL-1β are representative of three independent repeats. Statistical significance was determined by unpaired student’s t-test (H, I) or two-way ANOVA (B, C, D, E, K, L, N, O, P, Q): *p < 0.05; **p < 0.01; ***p < 0.001; ****p < 0.0001.

To determine if the chimera-based strategy of inflammasome-bypass can be extended to a more subtle engineering approach, we examined the redesigned hCasp-4 quadruple mutant that displays ICE activity *in vitro*. Within *Casp-1/11^−/−^* cells expressing this LPS receptor, which contains only four amino acid substitutions, LPS electroporation stimulated IL-1β and LDH release (Figure 5J-L). In contrast and as expected, cells expressing WT hCasp-4 produced only LDH, but not IL-1β (Figure 5J-L). Consistent with IL-1β release by redesigned hCasp-4 being an inflammasome-independent process, IL-1β release was insensitive to the inhibitory activities of MCC950 (Figure 5K,L). To further verify that redesigned hCasp-4 could bypass NLRP3 inflammasomes and cleave and release IL-1β, we determined if these cells could respond to LPS + nigericin treatment. Consistent with the fact that these cells lack endogenous mCasp-1 and mCasp-11, LPS + nigericin treatment was unable to promote IL-1β release (Figure S6F, G). Thus, IL-1β cleavage after LPS electroporation is driven by the intrinsic activities of redesigned hCasp-4, independent of inflammasomes.

We investigated whether this inflammasome-independent pathway can be engaged in response infection by Gram-negative bacteria. We focused on *Salmonella enterica* serovar Typhimurium, which activates inflammasomes by mCasp-11, as well as NLRP3 and NAIP-NLRC4 (Doerflinger et al., 2020). As described above, we eliminated NLRP3 activities from consideration in these studies, as the iBMDMs used are unresponsive to NLRP3 agonists. In addition, we used a flagellin-deficient strain of *Salmonella* (SL1344 *fliC/fljB*) (Wynosky-Dolfi et al., 2014) to diminish activation of the NAIP-NLRC4 inflammasome (Rauch et al., 2017). Consistent with this experimental setup focusing attention on LPS-induced activities, infection with SL1344 *fliC/fljB* induced LDH release in a hCasp-4 or mCasp-11 transgene-dependent manner (Figure 5N, O). Consistent with our findings with electroporated LPS, infection-induced IL-1β release was strongly enhanced within cells expressing redesigned caspases that contain LPS-binding and ICE activity (hCasp-4 quadruple mutant, hCARD4/cEnz or mCARD11/cEnz) (Figure 5P-S). In contrast. Cells expressing WT hCasp-4 or mCasp-11 were unable to release cleaved IL-1β after infection (Figure 5R, S). These collective data demonstrate the redesign of a normally inflammasome-dependent process into a one-protein signaling pathway that intrinsically links LPS detection to IL-1β cleavage and release.

### Cats naturally encode a one-protein signaling pathway that bypasses inflammasomes to link LPS detection with IL-1 cleavage and release

Our finding that the inflammasome pathway can be bypassed by the simple introduction of four mutations into hCasp-4 raises the question of whether similar caspases and pathways already exist in nature. To address this possibility, we performed a cell-based mini-screen, where we introduced Casp-4 homologues from species spanning a broad range of mammalian orders (namely mouse, cat, lemur, rabbit, sheep, horse, bat, manatee and wombat) into *Casp-1/11^−/−^* iBMDMs (Figure 6A). We then electroporated cells with LPS and assessed IL-1β release and pyroptosis. With the exception of the manatee and wombat genes, all Casp-4 homologues responded to LPS electroporation with the induction of pyroptosis (Figure 6B). The observation that manatee and wombat Casp-4 do not support LPS-induced pyroptosis is interesting, as these species are evolutionarily most distant from the rest of the animals in our panel. This finding hints that recognition of LPS might not be a primordial function of Casp-4, but was acquired after these species diverged. Another caspase that stands out, is the Casp-4 homologue from the cat (feline Casp-1/4b). Unlike all other caspases examined, *Casp-1/11^−/−^* iBMDMs expressing feline Casp-1/4b released large amounts of mature IL-1β as determined by ELISA and immunoblot (Figure 6C, D). As observed with our synthetically redesigned pathways, IL-1β release induced by feline Casp-1/4b was insensitive to MCC950 (Figure 6E, F). This finding suggests that IL-1β is processed and secreted upon activation of feline Casp-1/4b by LPS, without the need to assemble an inflammasome. Cats, but not dogs, therefore have the ability to mount an inflammatory response to cytosolic LPS, and they do so by employing a natural one-protein signaling pathway (Figure 6G).

**Figure 6:**
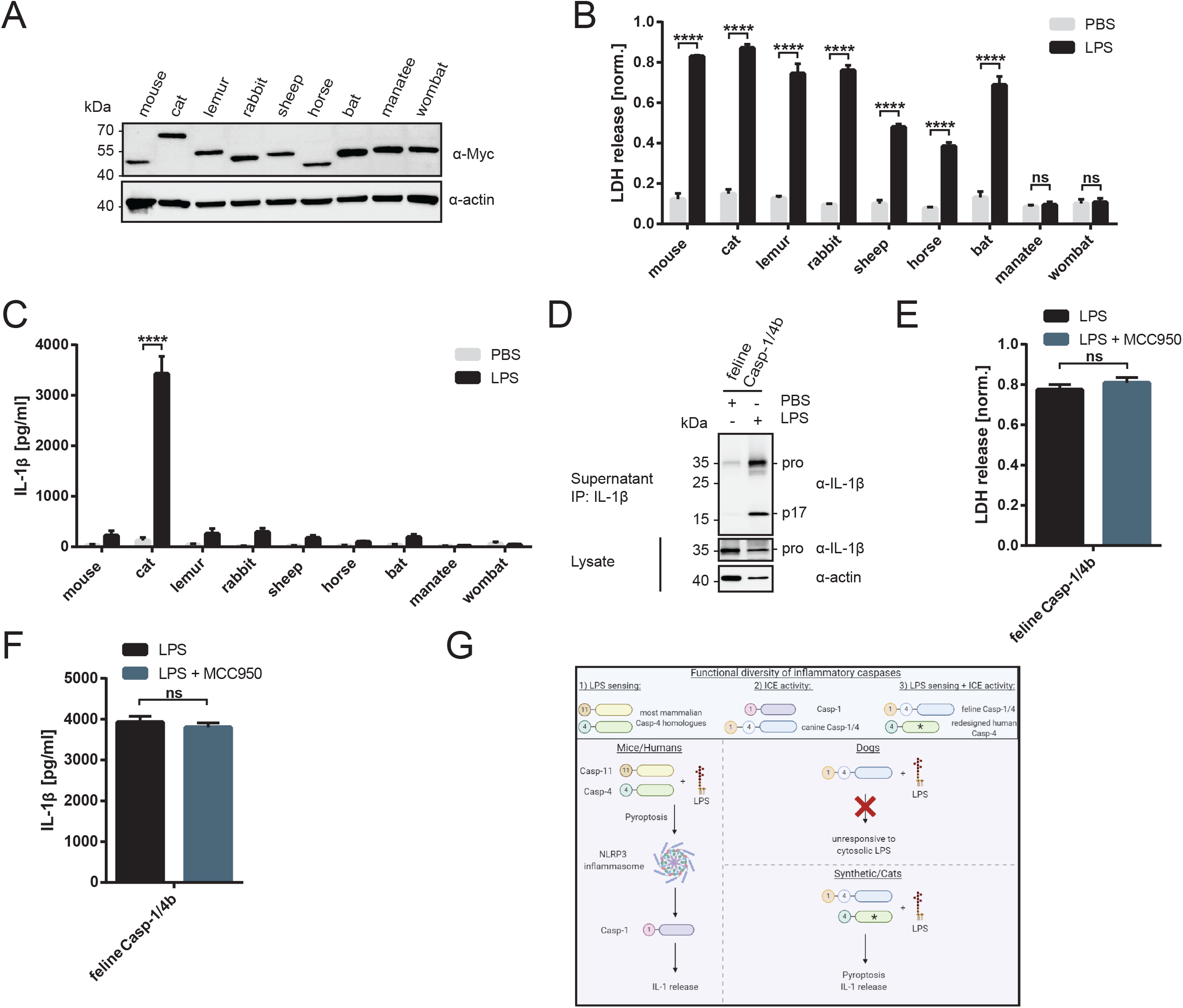
A natural one-protein signaling pathway that links LPS detection to IL-1β cleavage and release is present in cats. (A) *Casp-1/11^−/−^* iBMDMs were reconstituted by retroviral transduction with Casp-4 homologues from following mammalian species: mouse (*Mus musculus*), cat (*Felis catus*), lemur (*Microcebus murinus*), rabbit (*Oryctolagus cuniculus*), sheep (*Ovis aries*), horse (*Equus caballus*), bat (*Pteropus alecto*), manatee (*Trichechus manatus latirostis*), wombat (*Wombatus ursinus*). All constructs were detected in whole-cell lysates by immunoblot via an N-terminal Myc-tag. (B, C) *Casp-1/11^−/−^* iBMDMs reconstituted with the casp-4 homolog from the indicated species were primed for 4 h with LPS (1 μg/ml) before electroporation of LPS or PBS as negative control. LDH release and IL-1β levels in cell-free supernatants were quantified after 3 h using a colorimetric assay and ELISA, respectively. (D) *Casp-1/11^−/−^* iBMDMs reconstituted with feline casp-1/4 were primed for 4 h with LPS (1 μg/ml), followed by delivery of LPS (or PBS) into the cytosol via electroporation. Immunoprecipitated IL-1β from cell-free supernatants and cell-associated pro-IL-1β in whole-cell lysates were analyzed by immunoblotting after 3 h. (E, F) *Casp-1/11^−/−^* iBMDMs reconstituted with feline casp-1/4b were primed for 4 h with LPS (1 μg/ml), followed by delivery of LPS into the cytosol via electroporation. The NLRP3 inhibitor MCC950 (10 μM) was present in the cell culture media after electroporation. LDH release and IL-1β levels in cell-free supernatants were quantified after 3 h using a colorimetric assay and ELISA, respectively. (G) Model of synthetic and natural LPS-induced pyroptosis pathways in different mammals. Data are represented as mean ± SEM of three independent experiments. Immunoblots of immunoprecipitated IL-1β are representative of three independent repeats. Statistical significance was determined by two-way ANOVA (B+C) or unpaired student’s t-test (E+F): ****p < 0.0001.

## Discussion

In this study, we discovered that bioinformatic alignments failed to accurately predict the functions of cCasp-1/4 enzymes. Whereas one of the CARDs in cCasp-1/4b is most similar to LPS binding CARDs, cCasp-1/4b does not recognize LPS and canine cells do not respond to cytosolic LPS. Similarly, the enzymatic domains within both cCasp-1/4 isoforms closely resemble that from hCasp-4, yet cCasp-1/4 proteins display ICE activity *in vitro* and within cells. cCasp-1/4a and cCasp-1/4b also have the ability to facilitate pyroptosis through cleavage of GSDMD. Based on this evidence, we propose that canine Casp-1/4 proteins represent functional homologues of Casp-1. This disconnect between bioinformatic predictions based on homology and true biochemical functions is reminiscent of comparisons made between the Toll pathway regulators in *Drosophila* and mice. Upon TLR activation in mice, the protein TIRAP recruits MyD88 to assemble a SMOC known as the myddosome to stimulate immunity (Bonham et al., 2014). Flies contain a gene known as dMyD88, which had been considered the homologue of MyD88, based on sequence similarity. Biochemical and genetic analysis, however, revealed that dMyD88 is not the functional homologue of MyD88. dMyD88 is the functional homologue of TIRAP, which recruits the actual MyD88 homologue Tube to *Drosophila* Toll (Marek and Kagan, 2012). We speculate that these bioinformatic—function disconnects may become more common, as it is increasingly recognized that each species has tailored its immune networks in different manners (Daugherty and Malik, 2012).

A noteworthy outcome of our study of canine innate immunity is that it provided unexpected and novel insight into human and mouse biology. Indeed, carnivoran caspases proved to be excellent tools to identify mechanisms of substrate selection by Casp-1, a gene not even present in this lineage of animals. Proteomic and biochemical studies established that Casp-1 can cleave a broader range of substrates than mCasp-11 and hCasp-4, but the molecular determinants of these differential specificities remained largely unclear (Agard et al., 2010; Ramirez et al., 2018). Our analysis revealed that the ability of cCasp-1/4 and Casp-1 to cleave pro-IL-1β is conferred by a small set of charged amino acids located in proximity to the active site. By mutating select residues in hCasp-4 or mCasp-1, we successfully swapped the activities of these enzymes. For example, we generated a hCasp-4 mutant with substrate specificity similar to Casp-1 and a mCasp-1 mutant with substrate specificity similar to mCasp-11. These findings highlight how ICE activity is conferred to select caspases, which is a process distinct from that which facilitates GSDMD cleavage.

Recognition and cleavage of GSDMD is mediated by hydrophobic interactions between inflammatory caspases and the C-terminal domain of GSDMD (Liu et al., 2020; Wang et al., 2020). In contrast to these hydrophobic interactions that mediate GSDMD cleavage, our data indicate that pro-IL-1β cleavage is mediated by distinct electrostatic interactions. This idea is also supported by the fact that pro-IL-1β itself, especially the N-terminal pro-domain, is a highly charged molecule with distinct clusters of acidic residues surrounding its internal caspase cleavage sites. A notable aspect of the amino acids we identified is that they appear to specifically affect ICE activity, rather than overall catalytic activity of the caspase. In contrast, a recent study identified mutations that are conserved in some species of bats that dampen the inflammatory capacity of Casp-1 (Goh et al., 2020). However, these mutations most likely inhibit Casp-1 homodimerization, thereby interfering not only with IL-1β processing, but also cleavage of GSDMD and Casp-1 itself.

The knowledge gained from the study of canine caspases not only identified molecular determinants of ICE activity within Casp-1, but also illustrated the plasticity of network design. Our data indicate a direct correlation between the ability of a caspase to cleave pro-IL-1β *in vitro* and the degree to which it promotes IL-1β secretion from a cell, independent of the cell biological path that leads to its activation. While we found that recruitment of the enzymatic domain of hCasp-4 or mCasp-11 into an inflammasome does not promote IL-1β release, a caspase with ICE activity will cause IL-1β release, no matter if its activation was inflammasome-dependent or initiated by an interaction with LPS. We exploited this relationship to design synthetic one-protein signaling pathways that intrinsically link LPS detection to IL-1β cleavage and release via GSDMD, bypassing the need for an inflammasome. Strikingly, we discovered that similar pathways naturally exist in cats. This simplified means of signal propagation contrasts with the complex pathway architecture that is commonly observed in innate immunity. While complex pathways allow for intricate ways of regulation of signal transduction (Lee and Yaffe, 2016), multistep processes might also offer more opportunities for pathogens to develop immune evasive strategies. The unique synthetic and natural one-protein signaling pathways described in this study reveal unexpected network-design flexibility in nature, which will provide tools to investigate the potential cost and benefits of different types of innate immune pathway design.

## Acknowledgements

We thank Igor Brodsky for providing the flagellin-deficient *Salmonella* strain and Hao Wu for providing purified GSDMD. Cell sorting was performed at and with assistance by staff from the Harvard Immunology Flow Cytometry Core Facility or the Harvard Digestive Disease Center Flow Cytometry Core Facility. We thank all members for the Kagan lab for helpful input and discussions. This work was supported by NIH grants AI133524, AI093589, AI116550 and P30DK34854 to J.C.K. J.C.K. holds an Investigators in the Pathogenesis of Infectious Disease Award from the Burroughs Wellcome Fund. P.D. is supported by a fellowship by the Boehringer Ingelheim Fonds.

## Author Contributions

P.D. designed and performed experiments. A.C. generated and provided essential reagents.

J.C.K supervised all research. All authors discussed results and commented on the manuscript.

## Supplementary figure legends

**Figure S1: cCasp-1/4 proteins functionally complement for Casp-1 in response to various inflammasome stimuli.**

(A) Brightfield microscopy images of primary monocytes from canine whole blood on the day of isolation (Day 0) and after 6 days of differentiation in the presence of 50 ng/ml M-CSF to generate monocyte-derived macrophages (MDMs).

(B) Canine primary MDMs were primed with LPS (1 μg/ml) for 4 h or left unprimed before stimulation with nigericin (10 μM). Cells were pre-treated with indicated inhibitors for 30 min and inhibitors were co-administered during nigericin treatment. Cellular ATP levels were determined by Celltiter-Glo assay after 3 h.

(C, D) WT iBMDMs or *Casp-1/11^−/−^* iBMDMs reconstituted with the indicated caspase were primed for 4 h with LPS (1 μg/ml) or left unprimed before transfection with 5 μg/ml of poly(dA:dT) (DNA). Cell lysis and IL-1β in cell culture supernatants were quantified 6 h post-transfection by LDH assay and ELISA, respectively.

(E) Schematic representation of canine caspase construct lacking its first N-terminal CARD (cCasp-1/4bΔCARD) in comparison to WT cCasp-1/4 isoforms.

(F) Immunoblot analysis of whole-cell lysates of *Casp-1/11^−/−^* iBMDMs reconstituted with cCasp-1/4bΔCARD or WT cCasp-1/4 proteins. All constructs carry an N-terminal Myc-tag for detection.

(G, H) *Casp-1/11^−/−^* iBMDMs reconstituted with the indicated caspase were primed for 4 h with LPS (1 μg/ml) or left unprimed before stimulation with nigericin (10 μM). Cell lysis and IL-1β in cell culture supernatants were quantified after 3 h by LDH assay and ELISA, respectively.

Data are represented as mean ± SEM of at least three independent experiments. Statistical significance was determined by one-way ANOVA (B) or two-way ANOVA (C,D,G,H): *p < 0.05; ****p < 0.0001.

**Figure S2: Canine myeloid cells are unresponsive to cytosolic, but not extracellular LPS.**

(A) *Casp-1/11^−/−^* iBMDMs expressing mCasp-11 or cCasp-1/4bΔCARD were primed for 4 h with LPS (1 μg/ml) before electroporation of LPS or PBS as negative control. LDH in cell-free supernatants was quantified after 3 h using a colorimetric assay.

(B-D) Canine primary MDMs were primed with 1 μg/ml of LPS for 4 h. Subsequently, 1 μg of LPS was delivered into the cytosol of the cells by electroporation at a voltage of 1700 V. PBS electroporations served as negative controls.

(B) PI fluorescence intensity as a measure of plasma membrane rupture or pore-formation was quantified 3 h after electroporation.

(C) Cellular ATP levels were determined by Celltiter-Glo assay 3 h post-electroporation.

(D) Quantification of released IL-1β from cell-free supernatants was performed by ELISA after 3 h.

(E) Canine primary monocytes were stimulated with 1 μg/ml of extracellular LPS for 7 h or left untreated and IL-1β in cell-free supernatants was quantified by ELISA.

(F) Canine primary monocytes were stimulated with 1 μg/ml of extracellular LPS for 7 h or left untreated. IL-1β in the cell culture media was immunoprecipitated and analyzed by immunoblot.

Data are represented as mean ± SEM of three independent experiments. Immunoblots show representative result of three independent repeats Statistical significance was determined by unpaired student’s t-test (B-E) or two-way ANOVA (A): ***p < 0.001; ****p < 0.0001.

**Figure S3: Specific mutations in the catalytic domain of cCasp-1/4 reduce ICE activity, but not GSDMD cleaving activity.**

(A-C) 50 nM of recombinant murine pro-IL-1β was mixed with two-fold serial dilutions of chimeric caspase catalytic domains and incubated for 30 min at 37 °C. Proteins were then separated by SDS-PAGE and processing of pro-IL-1β was analyzed by immunoblotting. Catalytic efficiencies (k_cat_/K_m_) were calculated based on disappearance of band corresponding to pro-form of IL-1β and represented in comparison to WT caspases.

(D) Sequences alignment of enzymatic domains of select carnivoran Casp-1/4 enzymes. Sequences were aligned using the Clustal Omega online tool and visualized using ESPript 3.0. Amino acids of interest are indicated by red arrowheads.

(E-J) 50 nM of recombinant murine pro-IL-1β was mixed with two-fold serial dilutions of mutant cCasp-1/4 catalytic domains and incubated for 30 min at 37 °C. Proteins were then separated by SDS-PAGE and processing of pro-IL-1β was analyzed by immunoblotting.

(K, L) 50 nM of recombinant human GSDMD was mixed with two-fold serial dilutions of WT or mutant cCasp-1/4 catalytic domain and incubated for 30 min at 37 °C. Proteins were then separated by SDS-PAGE and processing of GSDMD was analyzed by immunoblotting.

Each data point (C) represents the result of one independent assay. Bars and error bars represent mean ± SEM of at least three independent experiments. Repeats of *in vitro* assays involved at least two independent caspase preparations. Immunoblots are representative of at least three independent repeats

**Figure S4: Specific mutations in the catalytic domain of hCasp-4 lead to increased pro-IL-1β cleavage *in vitro.***

(A-F) 50 nM of recombinant murine pro-IL-1β was mixed with two-fold serial dilutions of mutant hCasp-4 catalytic domains and incubated for 30 min at 37 °C. Proteins were then separated by SDS-PAGE and processing of pro-IL-1β was analyzed by immunoblotting.

(G, H) 50 nM of recombinant human GSDMD was mixed with two-fold serial dilutions of mutant hCasp-4 catalytic domains and incubated for 30 min at 37 °C. Proteins were then separated by SDS-PAGE and processing of pro-IL-1β was analyzed by immunoblotting.

(I) Homology model of cCasp-1/4 enzymatic domain (grey) bound to zVAD-FMK (pink) was generated using SWISS-MODEL. Amino acids of interest identified in this study are labeled in blue, main catalytic residue C285 is labeled in orange. Residue involved in recognition of GSDMD is yellow.

Repeats of *in vitro* assays involved at least two independent caspase preparations. Immunoblots show representative result of three independent repeats.

**Figure S5: H340 in Casp-1 is critical for the selection of IL-1β as a substrate.**

(A-D) 50 nM of recombinant murine pro-IL-1β was mixed with two-fold serial dilutions of mutant hCasp-4, mCasp-1 or cCasp-1/4 catalytic domains and incubated for 30 min at 37 °C. Proteins were then separated by SDS-PAGE and processing of pro-IL-1β was analyzed by immunoblotting. Catalytic efficiencies (k_cat_/K_m_) were calculated based on disappearance of band corresponding to pro-form of IL-1β and represented in comparison to those obtained for respective WT caspases.

(E) Sequence of alignment of Casp-4 and Casp-1 sequences from different mammalian species: human (*Homo sapiens*), mouse (*Mus musculus*), manatee (*Trichechus manatus latirostis*), cow (*Bos taurus*), pig (*Sus scrofa*), bat (*Myotis brandtii*) and beluga whale (*Delphinapterus leucas*).

(F, G) 50 nM of recombinant human GSDMD was mixed with two-fold serial dilutions of WT mCasp-1 or mCasp-1 carrying a H340D mutation and incubated for 30 min at 37 °C. Proteins were then separated by SDS-PAGE and processing of pro-IL-1β was analyzed by immunoblotting.

(H) 50 nM of recombinant murine pro-IL-1β was mixed with two-fold serial dilutions of purified mCasp-1 H340D catalytic domain and incubated for 30 min at 37 °C. Proteins were then separated by SDS-PAGE and processing of pro-IL-1β was analyzed by immunoblotting.

(I)*Casp-1/11^−/−^* iBMDMs reconstituted with WT mCasp-1 or mCasp-1 H340D mutant were primed for 4 h with LPS (1 μg/ml), then either left untreated or stimulated with nigericin (10 μM) for 3 h. Reactions were stopped by adding concentrated SDS loading buffer directly into the cell culture well to capture proteins in both the cell lysates and the cell culture supernatant. Proteins were separated by SDS-PAGE and processing of Casp-1 was analyzed by immunoblotting. Ticks mark location of full-length (FL) Casp-1 and cleavage fragments (p32, p20).

Each data point represents the result of one independent assay. Bars and error bars represent mean ± SEM of at least three independent experiments. Repeats of *in vitro* assays involved at least two independent caspase preparations. Immunoblots show representative result of three independent repeats. Statistical significance was determined by unpaired student’s t-test: *p < 0.05; **p < 0.01; ***p < 0.001; ****p < 0.0001.

**Figure S6: Design of NLRP3-independent signaling pathways that link IL-1β release to the detection of cytosolic LPS.**

(A) Schematic representation of chimeric caspases consisting of the enzymatic domain of cCasp-1/4 and the CARD of mCasp-11 and hCasp-4, respectively.

(B, C) *Casp-1/11^−/−^* iBMDMs expressing mCARD11/cEnz or hCARD4/cEnz were primed for 4 h with LPS (1 μg/ml), followed by delivery of LPS into the cytosol via electroporation. The NLRP3 inhibitor MCC950 (10 μM) was present in the cell culture media after electroporation. LDH release was quantified after 3 h using a colorimetric assay, respectively.

(D, E) WT iBMDMs were primed for 4 h with LPS (1 μg/ml), followed by delivery of LPS into the cytosol via electroporation. The NLRP3 inhibitor MCC950 (10 μM) was present in the cell culture media after electroporation. Cell lysis and IL-1β levels in cell-free supernatants were quantified after 3 h by LDH assay and ELISA, respectively.

(F, G) *Casp-1/11^−/−^* iBMDMs reconstituted with mCasp-1, indicated chimeras or hCasp-4 quadruple mutant were primed for 4 h with LPS (1 μg/ml) or left unprimed before treatment with nigericin (10 μM) for 3 h. LDH and IL-1β in cell culture supernatants were then quantified using a colorimetric assay and ELISA, respectively.

Data are represented as mean ± SEM of three independent experiments. Statistical significance was determined by two-way ANOVA (F+G) unpaired student’s t-test (B-E): *p < 0.05; **p < 0.01; ***p < 0.001; ****p < 0.0001.

## STAR Methods

### Contact for Reagent and Resource Sharing

Further information and requests for resources and reagents should be directed to and will be fulfilled by the Lead Contact, Jonathan C. Kagan

### Experimental Model and Subject Details

#### Cell lines

All cells were cultured in humidified incubators at 37 °C and 5% CO_2_. Immortalized bone marrow-derived macrophages (iBMDMs) from WT or *Casp-1/11^−/−^* mice were cultured in DMEM supplemented with 10% fetal bovine serum (FBS), Penicillin + Streptomycin, L-Glutamine and Sodium pyruvate, herafter referred to as complete DMEM (cDMEM). cDNA sequences for wildtype or chimeric caspase constructs of interest carrying an N-terminal Myc-tag were ordered as gBlocks (Integrated DNA technologies) and cloned into pMSCV IRES EGFP using *NotI* and *SalI* restriction sites. Point mutations were introduced using the Quikchange mutagenesis kit (Agilent) or the Q5 mutagenesis kit (NEB) according to manufacturer’s protocols. All constructs generated in this paper were sequence confirmed by Sanger sequencing. HEK293T cells were cultured in cDMEM and used as packaging cells for retroviral vectors. For the production of retroviral particles, 2.5×10^6^ HEK293T cells were seeded in a 10 cm cell culture dish. After overnight incubation at 37 °C, cells were transfected with 10 μg of pMSCV IRES EGFP encoding the protein of interest, 6 μg of pCL-ECO and 3 μg of pCMV-VSVG using Lipofectamine 2000 (ThermoFisher) according to the manufacturer’s instructions. After 18-24 h at 37 °C, media was changed to 6 ml of fresh cDMEM and virus containing supernatant was collected 24 h post media change. Supernatants were clarified from cellular debris by centrifugation (400 × g, 5 min) and filtered through a 0.45 μm PVDF syringe filter. ~2×10^6^ Caspase-1/11^−/−^ iBMDMs were transduced twice on two consecutive days in a 6-well plate by adding 4.5 ml of viral supernatant supplemented with Polybrene (1:2000; EMD Millipore) per well, followed by centrifugation for 1 h at 1250 × g and 30 °C. GFP^+^ cells were sorted twice on a FACSAria or FACSMelody cell corter (BD Biosciences) to obtain cell lines with stable and homogenous expression of the target protein. Transgene expression was confirmed by immunobloting using a rabbit anti-myc-tag or mouse anti-myc-tag primary antibody (both from Cell Signaling Technologies) at a 1:1000 dilution.

#### Differentiation and immortalization of bone marrow-derived macrophages

Bone marrow-derived macrophages were immortalized as described before with slight modifications (De Nardo et al., 2018). L929 fibroblasts secreting M-CSF were cultured in cDMEM and M-CSF containing supernatants were harvested, cleared by centrifugation (400 × g, 5 min) and passed through a 0.22 μm filter. CreJ2 cells were cultured in cDMEM and J2 retrovirus-containing supernatans were harvested, cleared by centrifugation (400 × g, 5 min) and passed through a 0.45 μm filter. Femurs from freshly sacrificed *Casp-1/11*^−/−^ mice (B6N.129S2-Casp1tm1Flv/J from Jackson Laboratories) were dissected and flushed with sterile PBS pH 7.4. Obtained cell suspension was passed through a 70 μm cell strainer before plating cells in non-tissue culture-treated 10 cm dishes (1 × 10^7^ cells per dish). Cells were cultured in cDMEM supplemented with 30% M-CSF-containing supernatants from L929 cells for 4 days. On day 4 and 6 post isolation, bone marrow cells were transduced with J2 virus by swapping media for immortalization media (30% L929 supernatant + 70% CreJ2 supernatant) for 24 h. Following day 7, cells were passaged in cDMEM supplemented with gradually decreasing doses of L929 conditioned media (starting at 30%) until they divided normally in unsupplemented cDMEM. When the concentration of supplemented L929 supernatant was below 5%, cells were cultured tissue culture-treated plates for better adherence.

#### Primary cell culture

Canine whole blood from healthy beagle dogs was purchased from BioIVT and processed within 24 h post blood draw. Canine primary cells were cultured in RPMI supplemented with 10% FBS, L-Glutamine, Sodium pyruvate and Penicillin + Streptomycin (referred to as complete RPMI) Whole blood was diluted at a 1:1 ratio with sterile PBS pH 7.4 + 2.5 mM EDTA before layering 30 ml of diluted blood over 15 ml of FiColl Paque PLUS density gradient media (GE Healthcare). Density gradient centrifugation was performed at 800 × g for 35 min at 20 °C. Total peripheral blood mononuclear cells were havested from the interphase and washed twice with wash buffer (PBS pH 7.4, 2.5 mM EDTA, 1% FBS). Red blood cells were lysed by resuspending the pellet in ACK lysis buffer and incubation for 5 - 10 min at RT. After a final washing step in wash buffer, total PBMCs were seeded in T75 cell culture flasks in 15 ml complete RPMI (40×10^6^ PBMCs per flask). After incubation for 2 h at 37 °C, non-adherent PBMCs were removed by three washes with pre-warmed, sterile PBS pH 7.4. Adherent cells (monocytes) were either used for experiments right away or cultured in complete RPMI supplemented with 50 ng/ml of recombinant human M-CSF (R&D Systems) for 5-7 days. Media was replenished with fresh complete RPMI containing M-CSF every 2 – 3 days.

### Method Details

#### Ligand and chemical reconstitution

*E. coli* LPS (serotype O:111 B4) was purchased from Enzo Biosciences as a ready-to-use stock solution of 1 mg/ml and used at a working concerntration of 1 μg/ml. Nigericin was bought from Invivogen, dissolved in 100% ethanol to 6.7 mM and used for NLRP3 stimulation at a concentration of 10 μM. Poly(dA:dT) was from Invivogen and dissolved to a stock concentration of 1 mg/ml. Poly(dA:dT) was transfected at a final concentration of 5 μg/ml. MCC950 and zVAD-FMK (both from Invivogen) were resuspended in sterile DMSO to a concentration of 20 mM and used at a final concentration of 10 μM and 20 μM, respectively. Disulfiram was purchased from Tocris Biosciences, dissolved in DMSO to generate a stock solution at 20 mM. During inhibition experiments, disulfiram was present at a concentration of 50 μM. To inhibit pyroptosis, cells were pre-treated with inhibitors for 30 min and the respective inhibitor was present during signal 2 of inflammasome stimulation. Propidium iodide solution (1 mg/ml) was purchased from MilliporeSigma. Recombinant human M-CSF (CHO expressed, carrier-free) was bought from R&D Systems and dissolved in sterile PBS pH 7.4 (stock concentration 50 μg/ml). YVAD-pNA was from Enzo Biosciences and the stock solution was prepared at 20 mM in DMSO.

#### Inflammasome activation, cell death assays and ELISAs

Inflammasome activation assays were routinely performed in a 96-well format. To activate the NLRP3 inflammasome, 1 × 10^5^ cells were seeded in duplicate wells in 100 μl of cDMEM and incubated for 1 – 2 h at 37 °C and 5% CO_2_ for cells to adhere to the plate before adding 100 μl of cDMEM with or without 2 × LPS (1 μg/ml final concentration) for priming. After 4 h at 37 °C and 5% CO_2_, cells were stimulated with 10 μM Nigericin (Invivogen) in 200 μl cDMEM for 3 h. Cell lysis was routinely quantfied employing the lactate dehydrogenase (LDH) release assay using the CyQuant LDH cytotoxicity assay kit (Thermo Fisher). Briefly, 50 μl of cell-free supernatants were mixed with 50 μl of LDH assay buffer and incubated for 15 – 30 min at RT in the dark. Absorbance at 490 nm and 680 nm was meaured on a Tecan Spark plate reader and signal was normalized to lysis controls. Since canine primary cells yielded low signal intensities in the LDH assay, incorporation PI in intracellular nucleic acids was used as an alternative readout to quantify membrane rupture/pore formation in experiments involving these cells. Cells were treated as decribed in a black 96-well plate with clear bottom and PI (Millipore Sigma) was added to the cell culture media at a 1:300 dilution for 30 min before the end of the stimulation. Plates were spun for 5 min at 400 × g to pellet all cells in the bottom plane of the plate and fluorescence was measured on a Tecan Spark device at an excitation wavelength of 530 nm and an emission wavelength of 617 nm. Cellular ATP levels as a measure of viability were determined using the Celltiter-Glo Luminescent Cell Viability kit (Promega). Supernatant was completely removed from the cells stimulated in a 96-well format before adding 30 μl of serum-free Opti-MEM (Thermo Fisher) and 30 μl of Celltiter-Glo substrate solution per well. After incubation for 2-5 min at RT while shaking, the mixture was transferred into a white 96-well plate and luminescence signal was quantified using a Tecan Spark plate reader. Release of IL-1β from murine or canine cells was assessed by ELISA using the IL1 beta mouse uncoated ELISA kit (Thermo Fisher) or the canine IL-1 beta/IL1F2 DuoSet kit (R&D Systems), respectively, according to manufacturer’s protocols. To activate the AIM2 inflammasome, 5 × 10^4^ cells were seeded in duplicate wells in 100 μl of cDMEM and incubated overnight at 37 °C and 5% CO_2_. After priming with 1 μg/ml of LPS for 4 h, cells were transfected with 5 μg/ml of Poly(dA:dT) (Inivivogen) using Lipofectamine 2000 in a total volume of 200 μl cDMEM per well. Cell culture supernatants for LDH assay and ELISA analysis were collected 6 h post transfection.

Processing of GSDMD and caspase-1 after inflammasome activation was analyzed by immunoblotting using a rabbit anti-GSDMD (Abcam) or mouse anti-murine caspase-1 p20 (Adipogen) primary antibodies; both diluted at 1:1000. 1 – 2 × 10^6^ iBMDMs of the cell line of interest were seeded in a 6-well plate and primed with 1 μg/ml of LPS for 3 – 4 h. In order to capture proteins present in both the cell lysate and the cell culture supernatant, signal 2 (10 μM of nigericin) was administered in 1.5 ml of serum-free Opti-MEM media for 3 h. Samples for immunoblotting were prepared by adding 375 μl of 5x SDS loading buffer directly to the well and heated to 65 °C for 10 min to fully denature proteins.

#### LPS electroporation

The Neon transfection system (ThermoFisher) was used to deliver bacterial LPS into the cytoplasm of cells. Cells were resuspended in R buffer at a density of 10 × 10^6^ cell/ml. LPS or sterile PBS pH 7.4 (as negative control) was mixed with the cell supension (1 μg of LPS or 1 ul of PBS per 1 × 10^6^ cells, respectively) before aspirating the cell suspension into the Neon electroporation pipette equipped with a 100 μl tip and performing electroporation with two pulses with a pulse width of 10 ms each and a voltage of 1400 V, unless stated otherwise. Cells were then dispensed into an appropriate volume of cell culture media without Penicillin + Streptomycin and seeded in 96-well or 6-well tissue culture plates. Cell death and IL-1β release was quantified 3h post electroporation by LDH or PI assay and/or CelltiterGlo assay, and ELISA, respectively, as descrived above.

#### Bacterial infections

*S. aureus* strain lacking the *oatA* gene (SA113 Δ*oatA*) was a kind gift from David Underhill (Cedars Sinai). Bacteria were streaked on TSA agar plates containing sheep blood (Thermo Fisher) and incubated overnight at 37 °C. Single colonies were picked to inoculate 5 ml of Todd-Hewitt Broth (Millipore Sigma) containing 25 μg/ml of kanamycin and grown up at 37 °C and 250 rpm for 24 h. Cultures were washed three times in PBS pH 7.4, and OD_600_ was determined before appropriately diluting bacteria in cDMEM without antibiotics. 1 × 10^5^ iBMDMs per well in a 96-well plate were primed with 1 μg/ml LPS or left untreated followed by infection with SA113 Δ*oatA* at an MOI of 30 in a total volume of 200 μl per well. Synchronized infection was facilitated by spinning the plates at 500 × g for 5 min right after addition of bacteria-containing media. After 1 h of infection, extracellular bacteria were killed by changing the media to cDMEM (with Pen/Strep) and supplemented with 50 μg/ml of gentamicin. Cell culture supernatants for LDH release assay and ELISA to quantify levels of IL-1β in the were collected 12 h post initial infection.

*Salmonella* strain deficient for flagellin (SL1344 *fliC/fljB*) was a kind gift from Igor Brodsky (University of Pennsylvania) and infections were performed as described before with minor modifications (Wynosky-Dolfi et al., 2014). Bacteria were streaked on LB agar plates containing 25 μg/ml kanamycin, grown overnight at 37 °C and plates were stored at 4 °C for later use. Overnight cultures (3 ml LB + 25 μg/ml kanamycin and 25 μg/ml chloramphenicol) were inoculated with a single bacterial colony and grown at 37 °C while shaking at 250 rpm. On the next morning, bacterial culture was diluted into high salt LB (3 ml LB + 100 μl of overnight culture + 78 μl of sterile 5 M NaCl) and incubated for another 3 h at 37 °C without shaking. Infections were performed in a 96-well (for LDH and ELISA analyses) or 12-well format (for IL-1β immunoprecipitations) using 1×10^5^ and 1×10^6^ cells per well, respectively. Cells were seeded in cDMEM without Pen/Strep and primed with 1 μg/ml of LPS for 3 h. Bacteria were washed three times with pre-warmed cDMEM without Pen/Strep and added to cells at an MOI of 100 in 200 μl (96-well) or 2 ml (12-well) of cDMEM without Pen/Strep. Synchronized infection was facilitated by spinning the plates at 500 × g for 5 min right after addition of bacteria-containing media and cells were incubated at 37 °C and 5 % CO_2_. After 1 h, gentamicin was added to a final concentration of 100 μg/ml to kill extracellular bacteria. Cell culture supernatants for downstream analyses were harvested at 4 h post-infection.

#### Immunoprecipitation of IL-1 from cell culture supernatants

IP of murine or human IL-1β from cell culture supernatants was performed as described before (Evavold et al., 2018). Supernatants from 0.5 – 1.0 × 10^6^ cells (cell number consistent within each individual experiment) stimulated as indicated were transferred into 5 ml FACS tubes and depleted of cells and debris by spinning at 400 × g for 5 min. Cell-free supernatants were transferred into new tubes and rotated overnight at 4 °C in the presence of 0.5 μg of biotinylated goat anti-murine IL-1β antibody (R&D Systems) and 20 μl neutravidin agarose beads (Thermo Fisher). Remaining cells in the well were lysed in 1x SDS loading dye and served as input control. Beads were subsequently washed three times with PBS pH 7.4 before eluting bound proteins in 50 μl of 1x SDS loading dye. Immunoprecipitated and cell-associated IL-1β was detected by immunoblotting using a rabbit anti-murine IL-1β antibody (Genetex). Cell-associated actin or tubulin was detected as a loading control using a mouse anti-tubulin antibody (DSHB hybridoma bank; 1:100 dilution) or a mouse anti-actin antibody from Sigma at a dilution of 1:5000. IP of canine IL-1β was performed equivalently using 0.18 μg of biotinylated canine IL-1β ELISA detection antibody (R&D Systems) as bait and rabbit anti-canine IL-1β antibody from BioRad for immunoblot detection. Unless stated otherwise, all primary antibodies were used at a concentration of 1:1000.

#### Recombinant protein expression and purification

DNA sequences encoding residues 131 - 404 of canine caspase-1/4 isoform a, residues 94 - 377 of human caspase-4 and residues 130 – 404 of murine caspase-1 were amplified by PCR using gBlocks of the full-length proteins as a template and cloned into a pET28A(+) vector with an N-terminal His6-tag using *BamHI* and *EcoRI* restriction sites. cDNA encoding full-length murine was synthesized by Integrated DNA Technologies and cloned equivalently. Chemically competent Rosetta (DE3) pLysS cells (EMD Millipore) were transformed with the plasmid of interest and plated on LB agar plates with 25 μg/ml of Kanamycin. Overnight pre-cultures inoculated with a single colony were grown at 30 °C and 250 rpm in 2x YT media containing 25 μg/ml kanamycin and 50 μg/ml chloramphenicol. Expression cultures of 500 – 1500 ml were inoculated with overnight cultures at a ratio of 1:100 and incubated at 37 °C and 250 rpm until the OD_600_ reached a value between 0.7 and 0.8. Following a cooling step on ice, protein expression was induced by adding IPTG to a final concentration of 0.25 mM and expression was allowed to proceed overnight at 18 °C. Bacterial pellets were harvested by centrifugation (5000 × g for 20 – 30 min at 4 °C) and stored at −20 °C if not immediately used for protein purification.

To purify recombinant proteins, bacterial pellets were resuspended in resuspension buffer (25 mM HEPES-NaOH pH 7.4, 150 mM NaCl, 10 mM Imidazole) and lysed by ultrasonication. Cell lysates were clarified by centrifugation (30 min, 20,000 × g, 4 °C) and passed through a 0.22 μm syringe filter before pouring them into a gravity flow column containing a bed of Ni-NTA agarose beads (Qiagen). Beads were washed with at least 10 bed volumes of wash buffer (25 mM HEPES pH 7.4, 400 mM NaCl, 25 mM Imidazole) and bound protein was eluted stepwise in resuspension buffer supplemented with 40 – 250 mM Imidazole. Fractions containing the protein of interest were pooled, buffer exchanged into storage buffer (25 mM HEPES-NaOH pH 7.4, 150 mM NaCl, 10% glycerol) using a PD-10 desalting column (GE Healthcare). Lastly, protein was concentrated by centrifugal ultrafiltration. Aliquots were snap-frozen in liquid nitrogen and stored at −80 °C for later use. Purity and integrity of the purified proteins were analyzed by SDS-PAGE followed by Instant*Blue* staining (Expedeon).

Recombinant human GSDMD was provided by Hao Wu (Harvard Medical School) and purified as described (Hu et al., 2020).

#### *In vitro* caspase cleavage assays

To analyze the cleavage of full-length protein substrates, two-fold dilution series of the indicated recombinant caspase was incubated with purified murine pro-IL-1β or human GSDMD at a final concentration of 50 nM in 40 μl of caspase assay buffer (10 mM PIPES pH 7.2, 10% sucrose, 10 mM DTT, 100 mM NaCl, 1 mM EDTA, 0.1% CHAPS) for 30 min at 37 °C. Reactions were stopped by adding 15 μl of 5x SDS loading dye and boiling at 65 °C for 10 min. Cleavage products were separated on a 12% SDS-PAGE gel and analyzed via immunoblotting using rabbit anti-murine IL-1β (Genetex) or rabbit anti-GSDMD (Cell Signaling) primary antibody at a 1:1000 dilution. Band intensities were densitometrically quantified using ImageJ to determine EC50 values and catalytic efficiencies were calculated as described before (Ramirez et al., 2018) using the following equation:

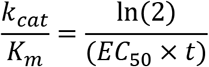

For peptide cleavage assays, recombinant mCasp-1, hCasp-4 or cCasp-1/4 were first diluted to a concentration of 100 nM in caspase assay buffer. To start the reaction, 20 μl of the diluted caspase were then mixed with 80 μl of a serial dilution of the chromogenic tetrapeptide substrate YVAD-pNA in the same buffer (final concentration of caspase was 20 nM in a total volume of 100 μl) in a clear 96-well plate. Absorbance at a wavelength of 405 nm was measured every 20 s for 20 min using a Tecan Spark plate reader with temperature control set to 37 °C. Substrate solution was pre-warmed to 37 °C before adding to the caspase to ensure homogenous assay conditions. Absorbance values were plotted in dependence of time and initial velocities were determined by performing linear fits of the resulting curves. A pNA standard curve was generated to transform absorbance values into molar concentrations. Initial velocities were plotted as a function of the substrate concentration and kinetic parameters (K_m_, v_max_, k_cat_) were determined by performing a fit according to the Michaelis-Menten equation in GraphPad Prism.

#### Sequence alignments and homology modelling

Mammalian caspase homologues were identified by NCBI BLASTp search using amino acid sequences of canine Casp-1/4a or human Casp-1 as search queries. Sequences of interest were aligned using the Clustal Omega Multiple Sequences Alignment tool and visualized in ESPript 3.0 (Madeira et al., 2019; Robert and Gouet, 2014).

To generate a homology model of the catalytic domain of cCasp-1/4, we utilized tools available on the SWISS-MODEL online server (Waterhouse et al., 2018). A crystal structure of the catalytic domain of human Casp-1 in complex with the inhibitor zVAD-FMK (sequence identity 57% and 63% for the p20 and p10 fragment, respectively) was used as an input template (pdb ID: 2H51; Datta et al., 2008). Obtained homology model was visualized in PyMol (Schroedinger, Inc.).

#### Quantification and Statistical Analysis

Statistical significance was determined by two-way ANOVA, one-way ANOVA, or unpaired student’s t-test as appropriate for each experimental dataset. Utilized statistical method is indicated in the respective figure legends. P-values below 0.05 were seen as statistically significant. All statistical analyses were performed using GraphPad Prism data analysis software. Data is representative of at least three independent repeats. Data with error bars are represented as mean ± standard error of the mean.

