## Supplementary figures and images for "Evolution-inspired dissection of caspase activities enables the redesign of caspase-4 into an LPS sensing interleukin-1 converting enzyme"

### Supplementary Figure 1

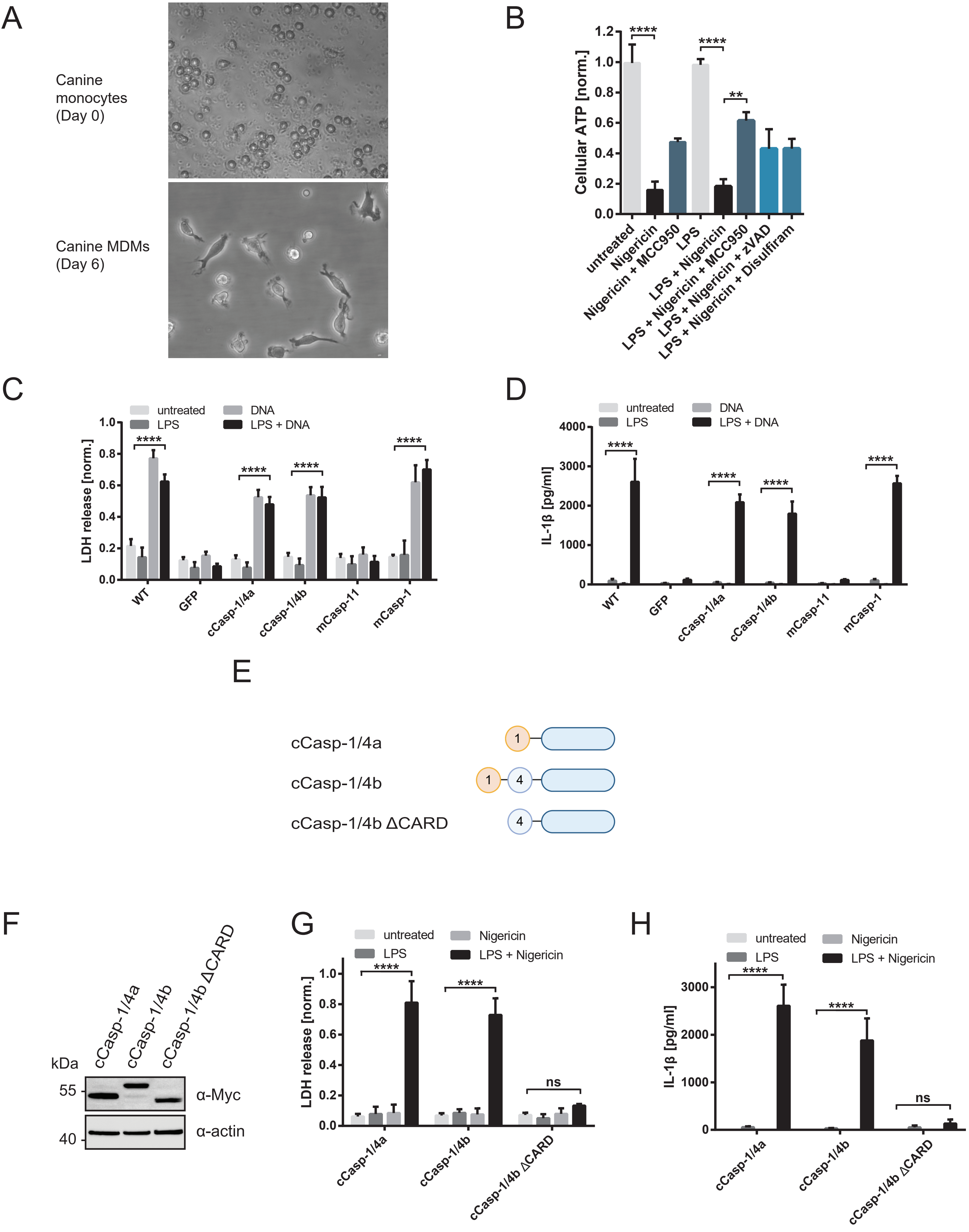

### Supplementary Figure 2

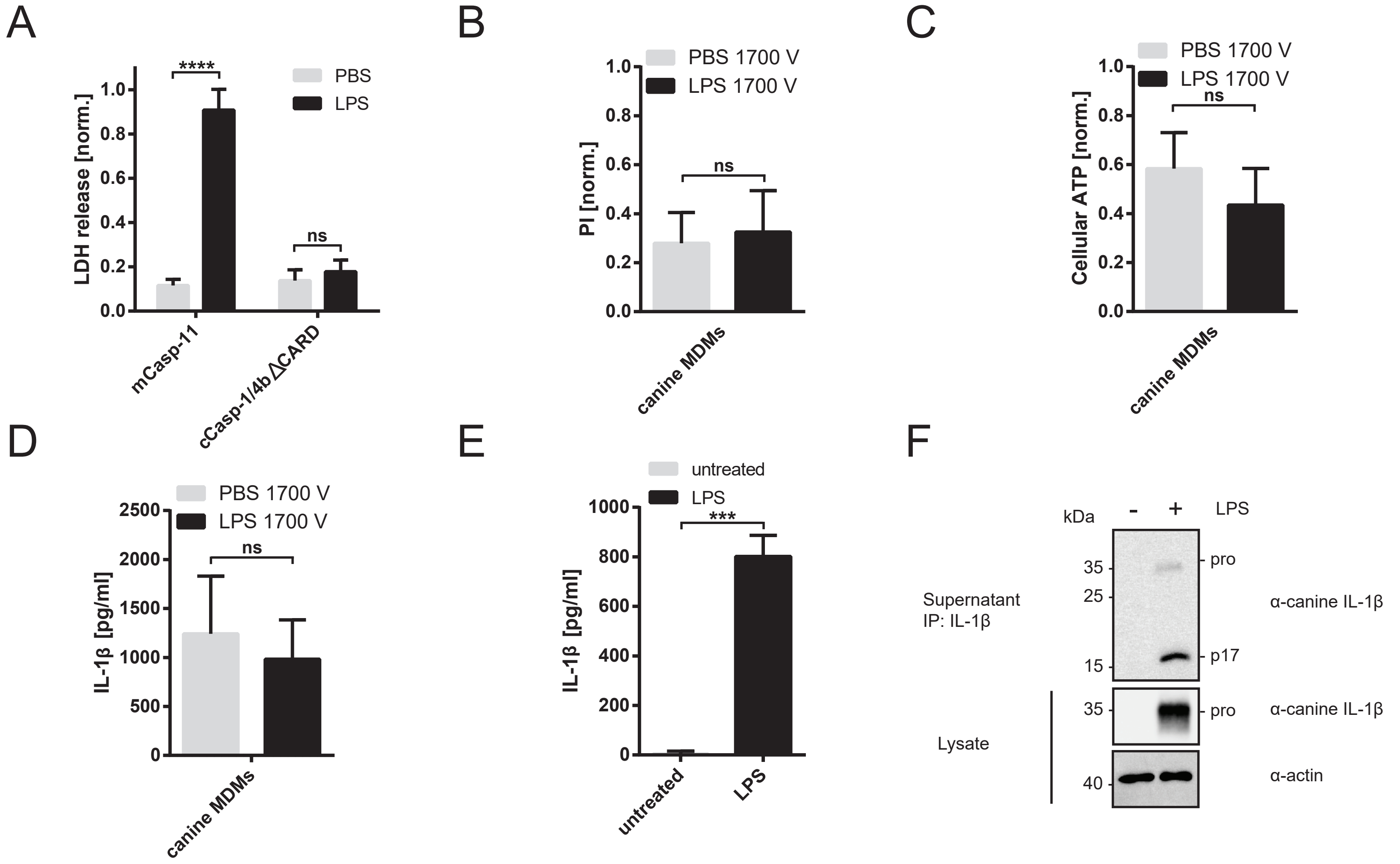

### Supplementary Figure 3

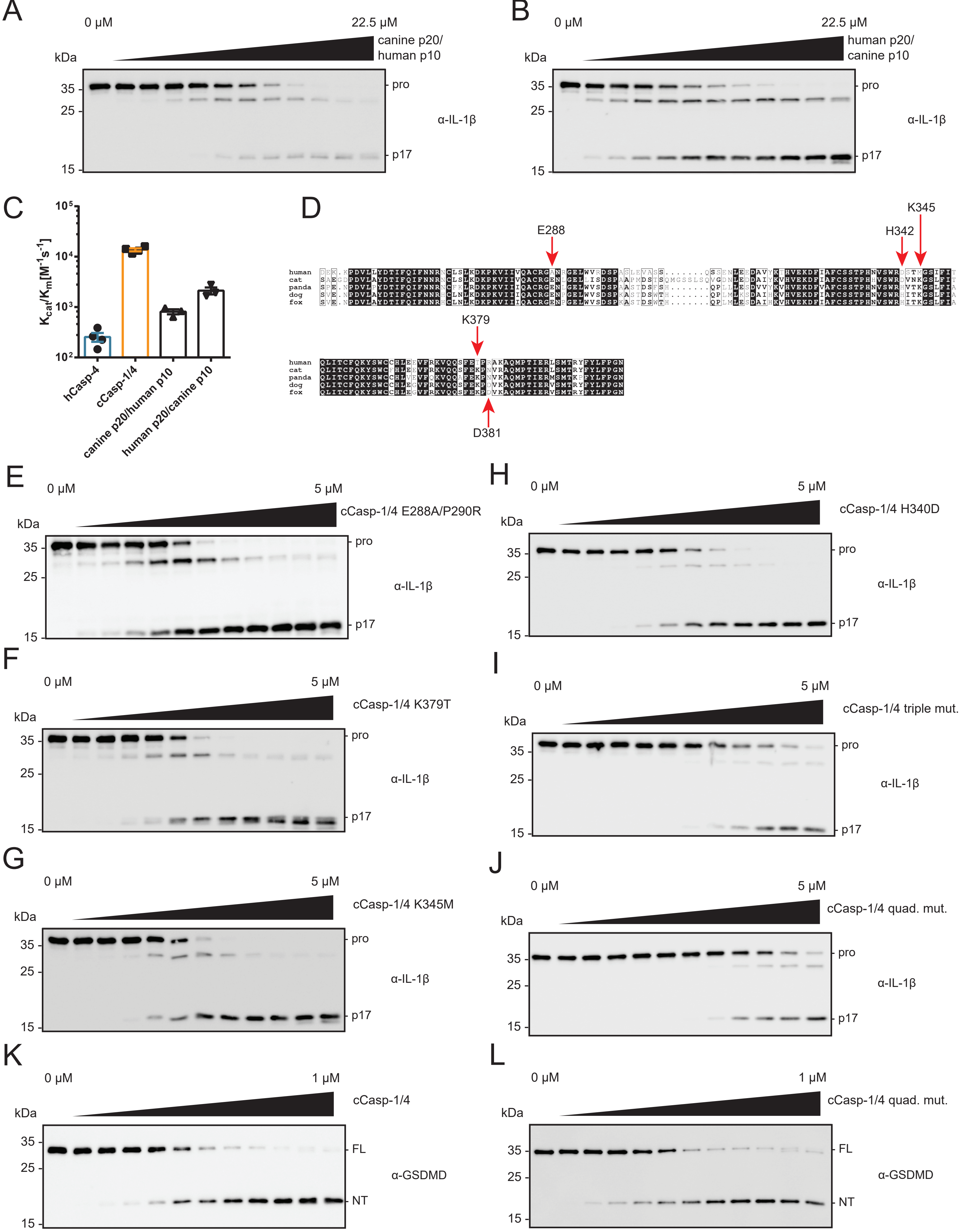

### Supplementary Figure 4

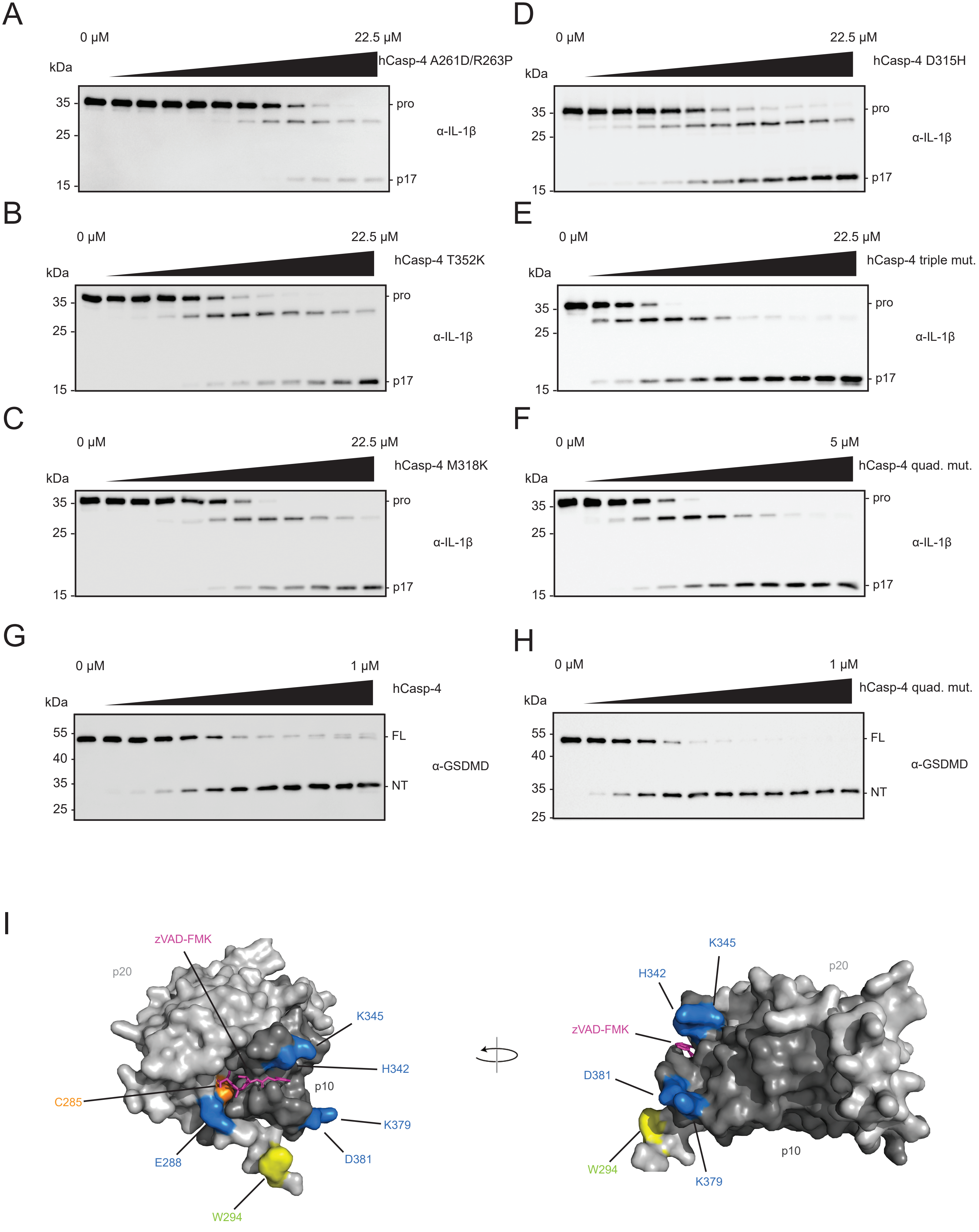

### Supplementary Figure 5

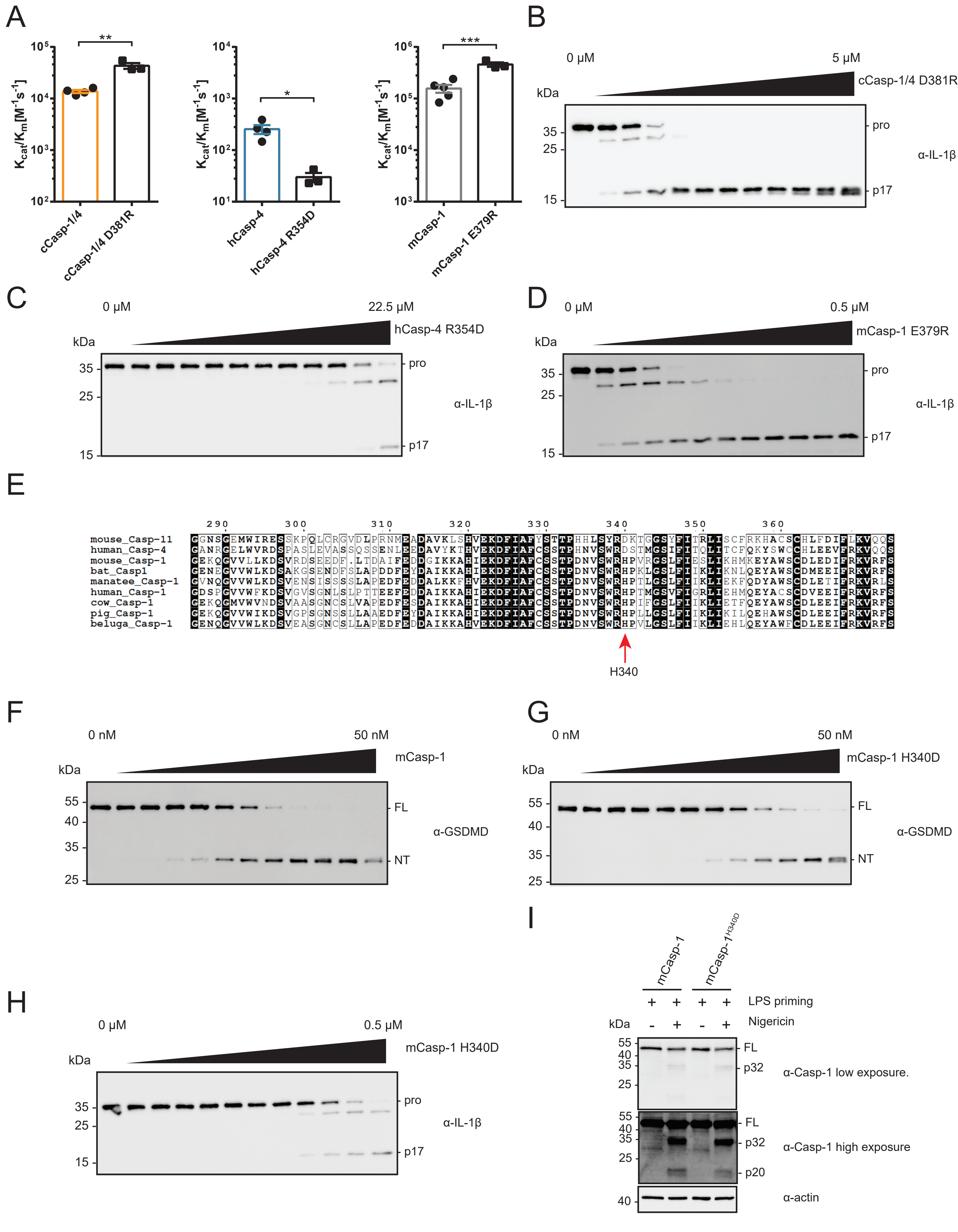

### Supplementary Figure 6

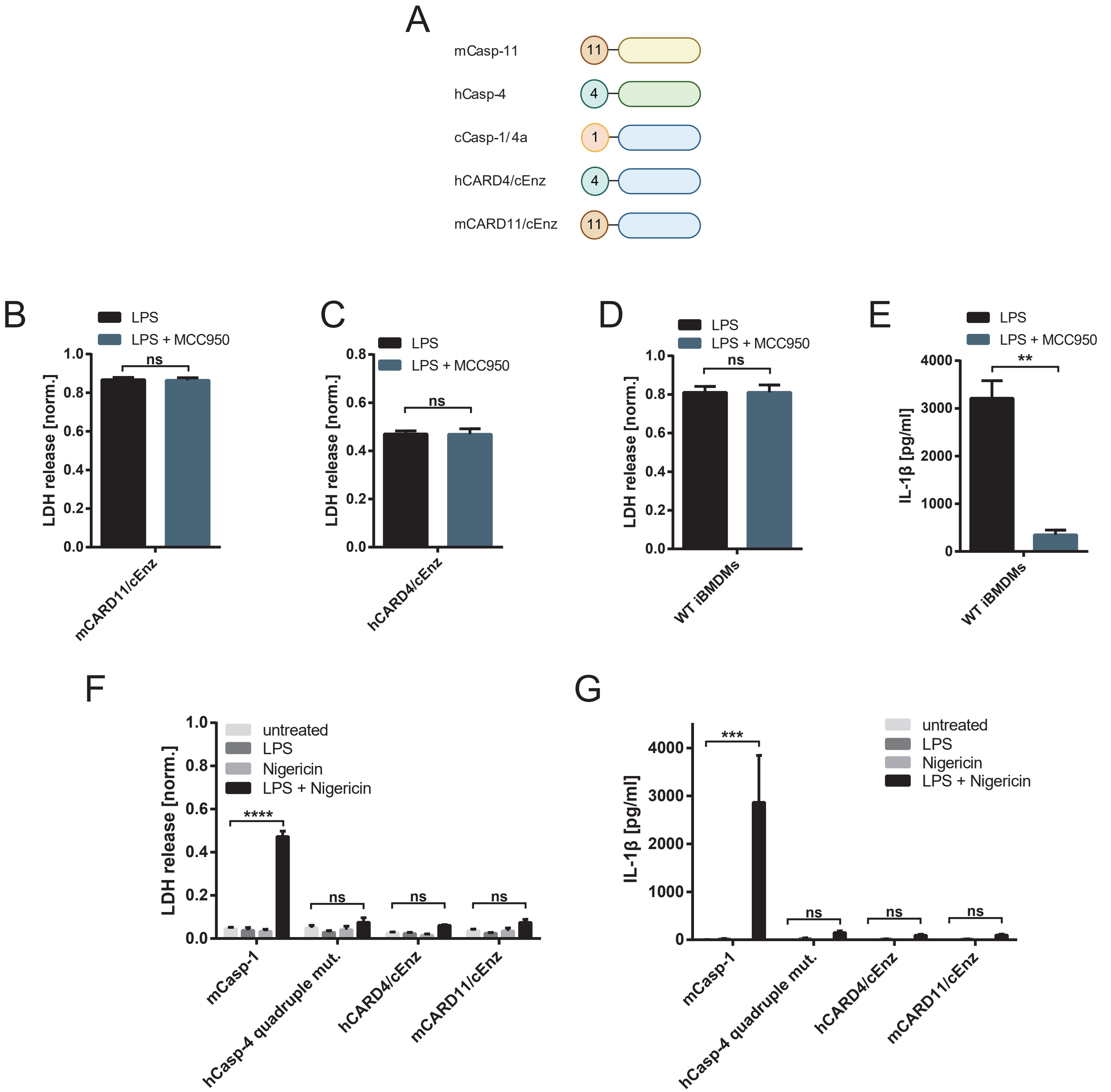
